# Eosinophils exert direct and indirect anti-tumorigenic effects in the development of esophageal squamous cell carcinoma

**DOI:** 10.1101/2023.06.01.543287

**Authors:** Justin Jacobse, Zaryab Aziz, Lili Sun, Jasmine Chaparro, Jennifer M. Pilat, Aaron Kwag, Matthew Buendia, Mae Wimbiscus, Motomi Nasu, Tsuyoshi Saito, Shinji Mine, Hajime Orita, Frank Revetta, Sarah P. Short, M. Kay Washington, Girish Hiremath, Michael K. Gibson, Lori Coburn, Tatsuki Koyama, Jeremy A. Goettel, Christopher S. Williams, Yash A. Choksi

**Author notes:** Corresponding author: Corresponding author, lead contact, Yash A. Choksi, M.D., Division of Gastroenterology, Hepatology, and Nutrition Vanderbilt University Medical Center, 2215 Garland Avenue 1075K MRB IV, Nashville TN, 37232.

## Abstract

**Background/Aims:** Eosinophils are present in several solid tumors and have context-dependent function. Our aim is to define the contribution of eosinophils in esophageal squamous cell carcinoma (ESCC), since their role in ESCC is unknown.

**Methods:** Eosinophils were enumerated in tissues from two ESCC cohorts. Mice were treated with 4-nitroquinolone-1-oxide (4-NQO) for 8 weeks to induce pre-cancer or 16 weeks to induce carcinoma. Eosinophil number was modified by monoclonal antibody to IL-5 (IL5mAb), recombinant IL-5 (rIL-5), or genetically with eosinophil-deficient (ΔdblGATA) mice or mice deficient in eosinophil chemoattractant eotaxin-1 (*Ccl11^-/-^*). Esophageal tissue and eosinophil specific RNA-sequencing was performed to understand eosinophil function. 3-D co-culturing of eosinophils with pre-cancer or cancer cells was done to ascertain direct effects of eosinophils.

**Results:** Activated eosinophils are present in higher numbers in early stage versus late stage ESCC. Mice treated with 4-NQO exhibit more esophageal eosinophils in pre-cancer versus cancer. Correspondingly, epithelial cell *Ccl11* expression is higher in mice with pre-cancer. Eosinophil depletion using three mouse models (*Ccl11^-/-^* mice, ΔdblGATA mice, IL5mAb treatment) all display exacerbated 4-NQO tumorigenesis. Conversely, treatment with rIL-5 increases esophageal eosinophilia and protects against pre-cancer and carcinoma. Tissue and eosinophil RNA-sequencing revealed eosinophils drive oxidative stress in pre-cancer. *In vitro* co-culturing of eosinophils with pre-cancer or cancer cells resulted in increased apoptosis in the presence of a degranulating agent, which is reversed with N-acetylcysteine, a reactive oxygen species (ROS) scavenger. ΔdblGATA mice exhibited increased CD4 T cell infiltration, IL-17, and enrichment of IL-17 pro-tumorigenic pathways.

**Conclusion:** Eosinophils likely protect against ESCC through ROS release during degranulation and suppression of IL-17.

## Introduction

In 2020, there were 604,100 new cases of esophageal cancer, 85% of which were ESCC.[1] More importantly, ESCC has an abysmal 5-year survival rate of less than 20%.[2, 3] In patients with localized disease a combination of chemoradiotherapy and surgery has led to a modest increase in median survival in recent years.[4, 5] However, patients who undergo esophagectomy have decreased quality of life, swallowing difficulties, malnutrition, and poor long-term survival;[6] thus, new therapies are needed for ESCC.

The tumor microenvironment has a central role in the growth of cancers.[7] The advent of immune checkpoint inhibitors to reactivate antitumor immunosurveillance [8] and clearance is a new treatment avenue for difficult to manage cancers. While these studies are focused mostly on T cells, myeloid cells are starting to be studied. [9, 10] Traditionally, studies centered on one of these myeloid cells, eosinophils, have focused on their role in allergic disease or helminth infection. However, recent studies have uncovered varying roles of eosinophils in cancers. Eosinophils are present in several solid tumors, including breast, colon, gastric, ovarian, and lung. [11] In colon cancer, they have been shown to be protective, [12] but in other cancers, such as cervical, [13, 14] Hodgkin’s lymphoma, [15, 16] and ovarian, [17] eosinophils have been shown to be pro-tumorigenic. However, relatively little is known about their contribution in esophageal cancer, though early studies suggest that tumor-associated tissue eosinophilia is associated with increased overall survival. [18–20] The role of eosinophils in different cancer types remains a topic of active research, as the interplay between eosinophils, other immune cells, and the tumor microenvironment is still incompletely understood. Understanding the role of eosinophils in cancer is important, as it could lead to new therapeutic strategies. In this study, we show that eosinophils infiltrate human ESCC to a greater extent in early versus late stages. We demonstrate that reduction in eosinophil number by different strategies exacerbates tumorigenesis. Conversely, treatment of mice with rIL-5 reduces esophageal tumor burden. Furthermore, we establish that eosinophils protect from murine ESCC through degranulation and subsequent release of ROS. Therapies targeting increased eosinophil recruitment to the esophagus may be useful in patients with ESCC.

## Results

### Eosinophils are present in human ESCC and specific to the tumor microenvironment

In order to determine whether eosinophilic presence in ESCC was tumor-specific, Eosinophil Peroxidase (EPX) IHC was performed on samples for which tumor-adjacent normal tissue was also available. EPX is an enzyme concentrated in granules inside eosinophils, and anti-EPX IHC is used as a histologic marker to identify eosinophils and detect the presence of degranulation. [12, 21]

Within the tumor, the number of EPX^+^ cells was higher as compared with adjacent normal (Figure 1A). Red arrows highlight the presence of degranulation, and extracellular deposition of eosinophil granule content which was seen in nearly all patient samples in which eosinophils were present. Patient demographic information is reported in Supplementary Table 2. Given that the esophagus does not normally have resident eosinophils, this suggested that recruitment of eosinophils was tumor-specific.

**Figure 1.**
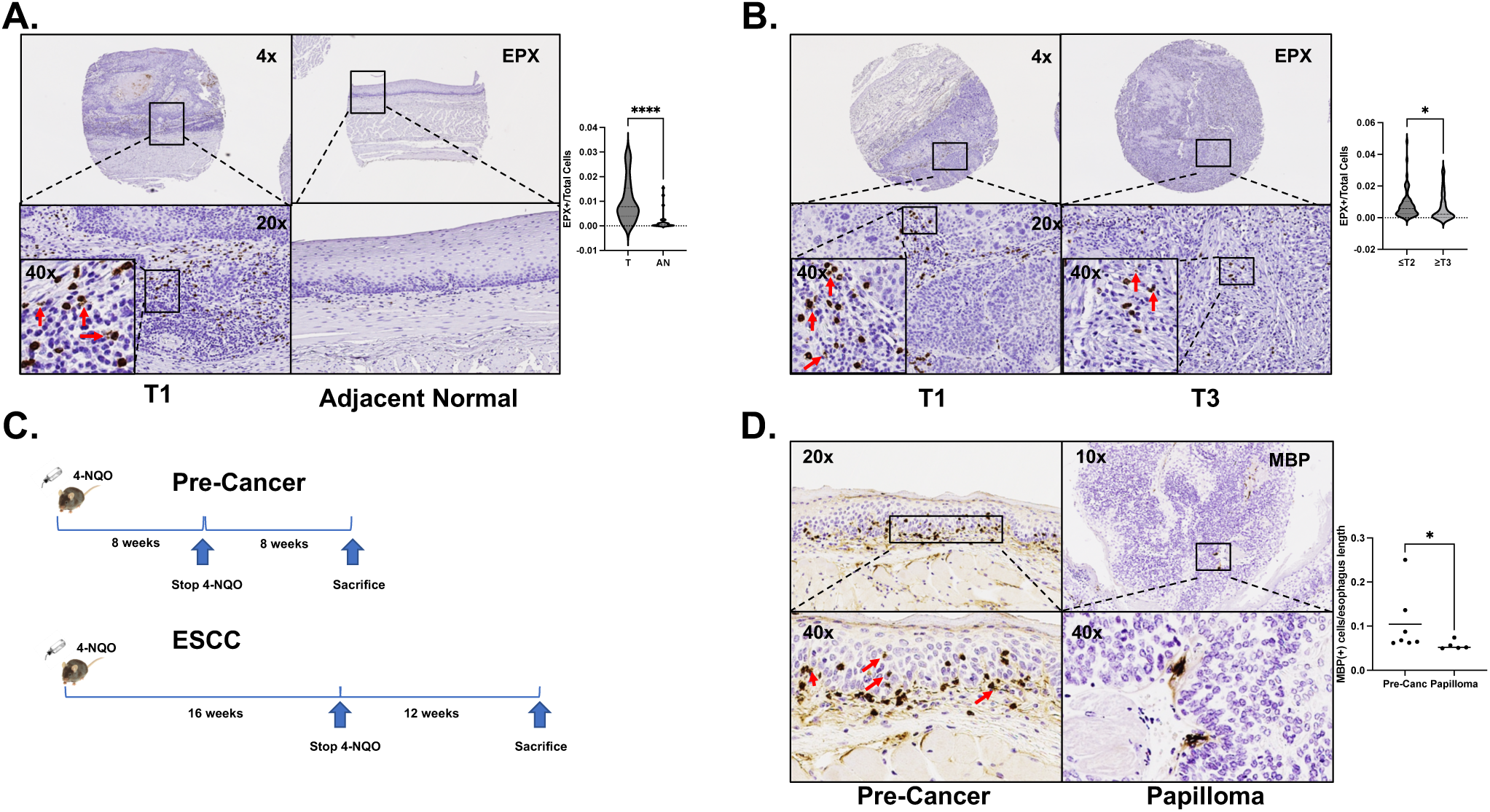
Eosinophils are present in greater numbers in patients with early stage Esophageal squamous cell cancer as compared with late stage. (A) There is a significantly greater ratio of EPX positive cells/total cells in human ESCC as compared with adjacent normal [0.0078 (0.0039, 0.015) vs 0.00048 (0.00011, 0.0011), P<0.0001, n=28)]. Red arrows point out degranulating eosinophils. (B) There is a significantly greater ratio of EPX positive cells/total cells in early stage ≤T2 human ESCC as compared with late stage ≥T3 ESCC [0.0053 (0.0023, 0.0095) vs 0.0026 (0.00068, 0.0097), P=0.04, n=38-67]. Data for A and B are represented with violin plots. Red arrows point out degranulating eosinophils. (C) Timelines for murine 4-NQO induced pre-cancer and carcinoma. In pre-cancer, 4-NQO is given via drinking water for 8 weeks followed by 8 weeks of propylene glycol vehicle. In carcinoma, 4-NQO is given via drinking water for 16 weeks followed by 12 weeks of propylene glycol vehicle. (D) There are significantly more MBP positive cells/length of pre-cancer in μm from mice treated with 4-NQO for 8 weeks as compared with papilloma/length of papilloma) from mice treated with 4-NQO for 16 weeks (0.10 ± 0.03 vs. 0.06 ± 0.005, P=0.03, n=5-7). Red arrows highlight degranulating eosinophils. All comparisons are reported as mean ± SEM and are using Mann-Whitney.

### Early stage ESCC is characterized by greater eosinophilic infiltration than late stage ESCC

We next compared eosinophil number in early stage ESCC (≤T2) with later stage ESCC (≥T3) and found that there were significantly more EPX^+^ cells in early stage cancers (Figure 1B). Again, the red arrows highlight the occurrence of degranulation, indicating the eosinophils present were activated. Patient demographic information for ≤T2 group as compared with ≥T3 group is listed in Supplementary Table 3. While the ≥T3 group was significantly younger, the difference in median age was 9 years and there was considerable overlap in the interquartile ranges. This observation suggests that eosinophils are being recruited to a greater degree early in tumorigenesis.

### Murine ESCC pre-cancer is characterized by greater eosinophilic infiltration as compared with carcinoma

The 4-NQO murine model of ESCC mimics the development of ESCC in humans. [22] 4-NQO contains similar carcinogens to those present in tobacco, a known risk factor for ESCC, [2, 3, 23] and causes DNA damage. [24–26]

Mice were treated with 4-NQO for 8 weeks followed by 8 weeks of vehicle or with 4-NQO for 16 weeks followed by 12 weeks of vehicle (Figure 1C). Mice treated with 4-NQO for 8 weeks had significantly more MBP^+^ cells in pre-cancerous areas as compared with papillomas or invasive carcinoma from mice treated with 4-NQO for 16 weeks (Figure 1D). To further investigate this, we also compared areas of papilloma or invasive carcinoma versus pre-cancerous areas in mice treated with 4-NQO for 16 weeks. There were significantly more MBP^+^ cells in pre-cancer areas as compared with papillomas or invasive carcinoma (Supplemental Figure 1). This is easily visualized in the 20x image. This finding provided evidence that murine development of ESCC in the 4-NQO model was similar to what occurs in human ESCC.

### Targeted eosinophil depletion exacerbates esophageal pre-cancer

Since we observed significantly greater eosinophilia in mice in pre-cancer versus in papillomas or invasive carcinoma, we hypothesized that eosinophils might protect from ESCC progression. Thus, we treated WT mice with a monoclonal antibody against IL-5 (IL5mAb), which blocks eosinophil differentiation, for the second eight weeks of the pre-cancer timeline (Figure 1C). Histology score was computed based on the severity of four characteristics: rete pegs, nuclear histologic abnormalities, papilloma, and invasive carcinoma. [27] Rete pegs and nuclear histologic abnormalities are common pre-cancer model findings, while papilloma and invasive carcinoma are common cancer model findings. Histology score showed significantly worse disease in mice treated with IL5mAb. The H&E highlights, in particular the 20x views, greater depth of rete pegs and the beginning of invasive disease, indicators of more advanced disease with eosinophil depletion (Figure 2A). While there was not a statistically significant difference in invasion, none of the control mice show invasive disease while 50% of the mice treated with IL5mAb did. We confirmed eosinophil depletion via MBP IHC in mice treated with IL5mAb (Figure 2B).

**Figure 2.**
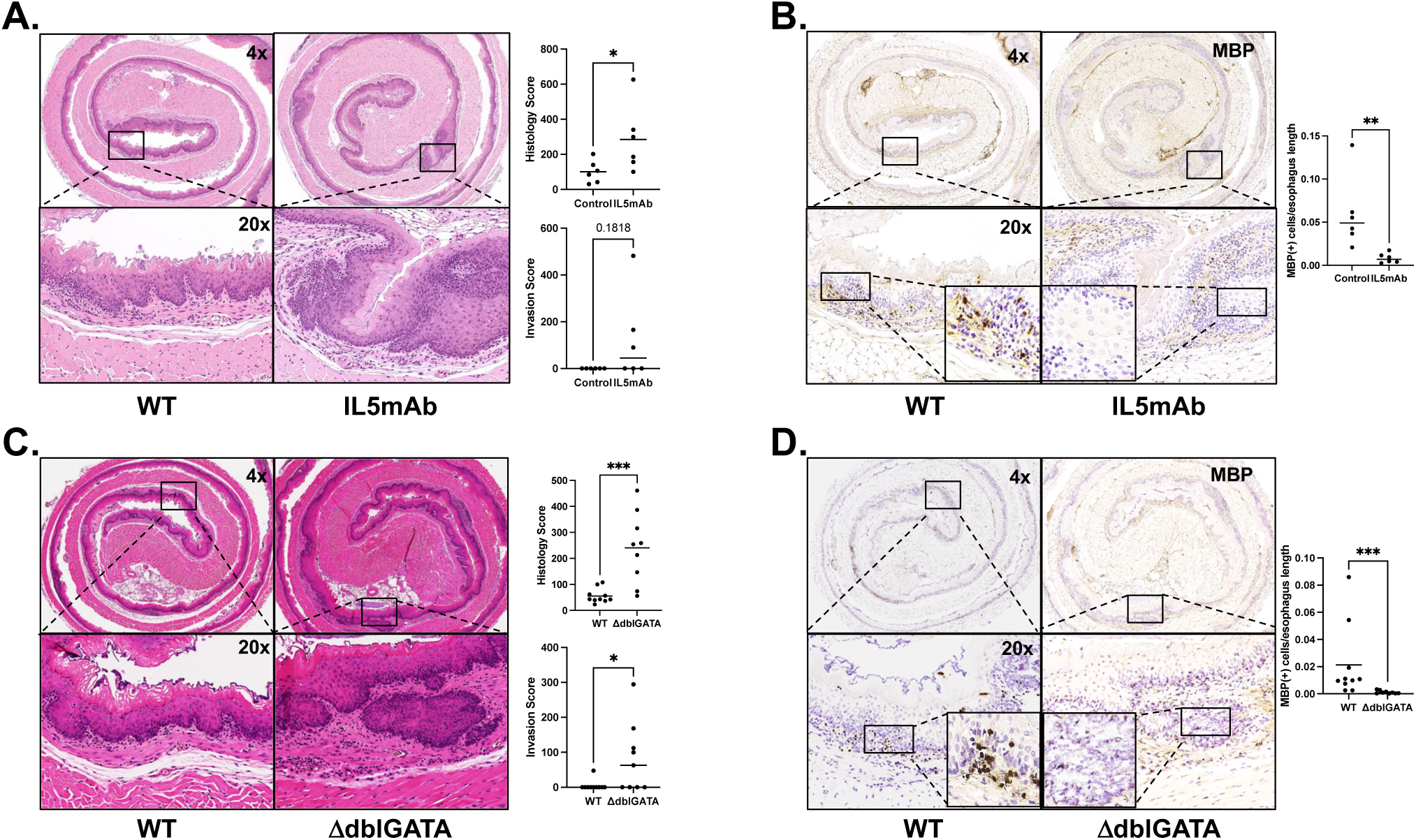
A reduction in eosinophils leads to significantly worse pre-cancer after 8 weeks of 4-NQO. (A) Mice treated with monoclonal antibody to IL-5 (IL5MAB) for the second eight weeks of the 4-NQO pre-cancer protocol had significantly increased histology score as compared with controls (284.6 ± 77.6 vs 100.8 ± 26.1, P=0.04, n=6). There is a trend towards an increased invasion parameter of the total score (122.7 ± 76.8 vs 0.0 ± 0.0, P=0.18, n=6), and all controls have a score of 0. (B) Mice treated with IL5MAB had significantly fewer MBP positive cells as compared with control (0.008314 ± 0.002308 vs 0.05944 ± 0.01702, P=0.002, n=6). Inserts on the lower two panels are taken at 40x magnification. (C) Eosinophil-deficient (ΔdblGATA) mice have significantly increased histology score as compared with controls in the pre-cancer protocol (240.7 ± 45.3 vs 55.6 ± 8.8, P<0.001, n=9-10). They demonstrate significantly increased invasion parameter of the total score (82.0 ± 33.4 vs 4.8 ± 4.8, n=9-10, P=0.02) as well. (D) There are significantly fewer MBP positive cells/esophageal length in ΔdblGATA mice as compared with WT controls (0.0011 ± 0.0004 vs 0.021 ± 0.0086, P<0.001, n=9-10). Inserts on lower two panels are taken at 40x magnification. All comparisons are reported as mean ± SEM and are using Mann-Whitney.

We next treated genetically eosinophil-deficient (ΔdblGATA) mice with 4-NQO in the precancer timeline. Confirming the phenotype observed with IL5mAb-mediated eosinophil depletion in WT mice, ΔdblGATA mice showed more advanced pre-cancer and invasion, which is highlighted in the 20x view (Figure 2C). In the 4x image, the rete pegs are apparent. MBP IHC confirmed the absence of eosinophils in ΔdblGATA mice (Figure 2D). As such, reducing eosinophil number either with IL5mAb or genetically with ΔdblGATA mice results in more advanced pre-cancer.

### Absence of eosinophils results in increased esophageal tumorigenesis

Since ΔdblGATA mice displayed more advanced pre-cancer as compared with controls, we hypothesized that ΔdblGATA mice would also develop more severe tumorigenesis in an ESCC cancer model (Figure 1C). As predicted, ΔdblGATA mice had significantly more esophageal papillomas than WT mice (Figure 3A). Additionally, there was a trend towards decreased survival in ΔdblGATA mice (Figure 3B), and due to this, the experiment was ended 4 weeks early to minimize the number of mice lost in the experiment. 4-NQO can induce oral tumors as well; however, no genotype specific differences in oral tumors was observed (Figure 3C). Thus, we hypothesized that this survival trend was due to esophageal tumorigenesis rather than oral tumorigenesis since there was no difference in tongue tumor number. Finally, consistent with the survival trend, we observed there was a significantly higher histology score and invasion score in ΔdblGATA mice (Figure 3D). Thus, ΔdblGATA mice demonstrate features of advanced precancer and cancer.

**Figure 3.**
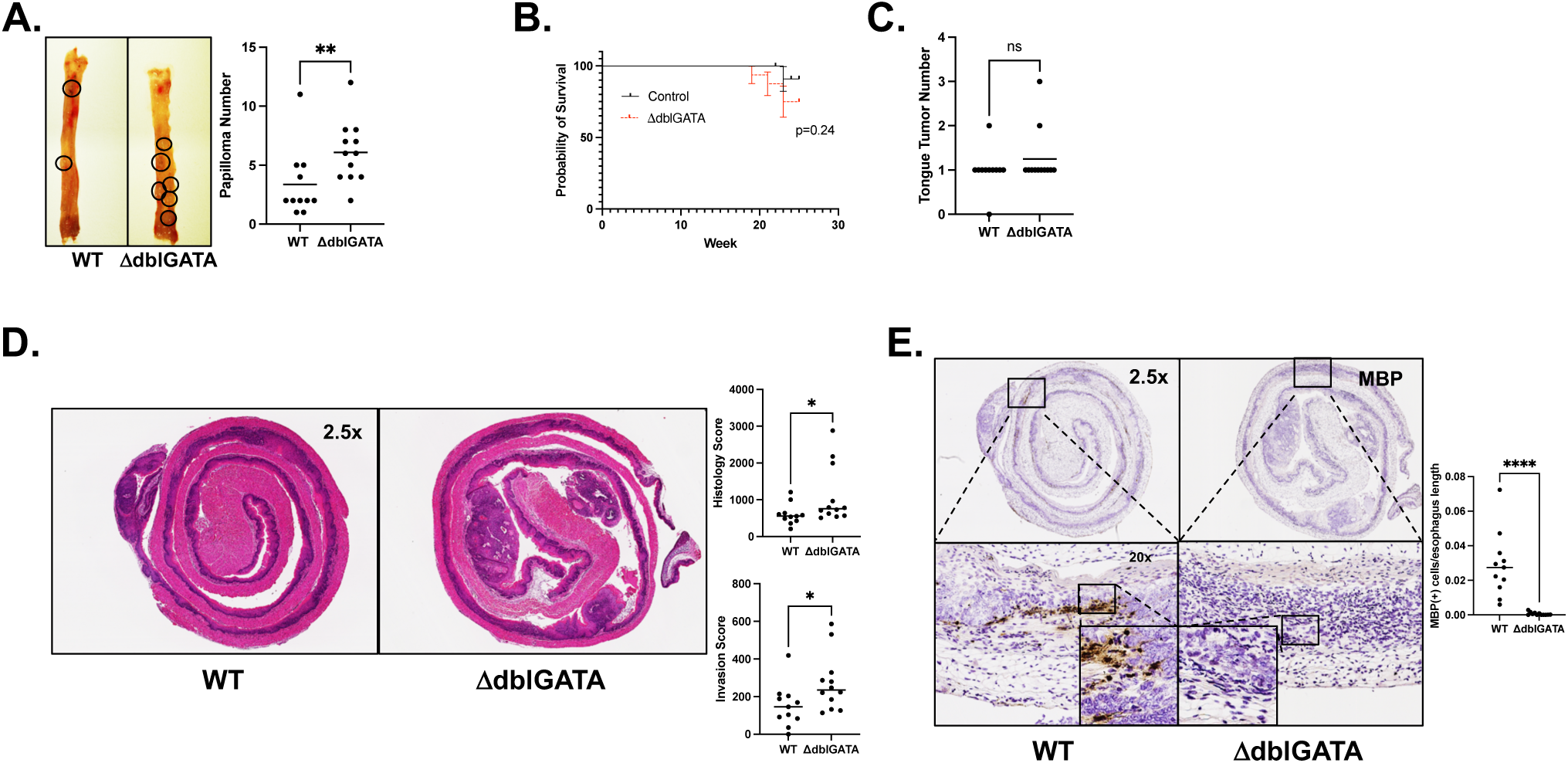
The absence of eosinophils leads to significantly worse carcinoma in a 4-NQO carcinoma timeline. (A) There are significantly more esophageal papillomas in ΔdblGATA as compared with WT control mice (6.08 ± 0.75 vs 3.36 ± 0.88, P<0.01, n=11-12). (B) There is a trend towards increased mortality for ΔdblGATA as compared with WT mice in the 4-NQO carcinoma model (P=0.24, n=12-16). (C) There is no difference in the number of tumors on the tongue between ΔdblGATA and WT mice. (D) ΔdblGATA have significantly increased histology score as compared with control mice in the 4-NQO carcinoma timeline (1107 ± 227.1 vs 601.7 ± 84.9, P=0.03, n=11-12). They demonstrate significantly increased invasion parameter of the total score (271.7 ± 43.4 vs 150.4 ± 33.9, P=0.02, n=11-12). (E) ΔdblGATA mice have significantly fewer MBP positive cells/esophagus length as compared with WT control mice (0.00062 ± 0.00026 vs 0.029 ± 0.0056, P<0.0001, n=11-12). All comparisons are reported as mean ± SEM and are using Mann-Whitney.

### Ccl11 is upregulated in pre-cancer

Since there were significantly more eosinophils in mice that were treated with 4-NQO in pre-cancer (8 weeks) versus cancer (16 weeks), we hypothesized that *Ccl11*, or *Eotaxin-1*, an eosinophil chemoattractant, which is expressed in the gastrointestinal tract,[28] may also be upregulated in pre-cancer. Thus, we queried a publicly available sc-RNAseq dataset [29] in which transcriptomic profiling was done at several time points throughout 4-NQO treatment. The authors of the study utilized a carcinoma model (16 weeks of 4-NQO, followed by 12 weeks vehicle) and sacrificed mice at these timepoints – 0 weeks, 12 weeks, 20 weeks, 22 weeks, 24 weeks, and 26 weeks timepoints after initiation of 4-NQO. They defined 0 weeks as normal (NOR), 12 weeks as inflammation (INF), 20 weeks as hyperplasia (HYP), 22 weeks as dysplasia (DYS), 24 weeks as carcinoma in situ (CIS), and 26 weeks as invasive carcinoma (ICA). After reviewing the histologic images, we determined that the 20-week hyperplasia timepoint was most similar to our “pre-cancer experiments” where mice were treated with 4-NQO for 8 weeks followed by 8 weeks vehicle. Thus, we compared *Ccl11* expression at 20 weeks versus the other timepoints. This analysis focused on epithelial cells and fibroblasts because *Ccl11* expression was very low in immune cells. In epithelial cells, *Ccl11* was significantly upregulated at 20 weeks as compared with 0, 12, 22, 24, and 26 weeks after the 4-NQO start date (Figure 4A). Additionally, *Ccl11* was significantly upregulated at 20 weeks in fibroblasts as compared with 0 and 22 weeks after initiation of 4-NQO but unchanged at 12, 24, or 26 weeks (Supplemental Figure 2).

**Figure 4.**
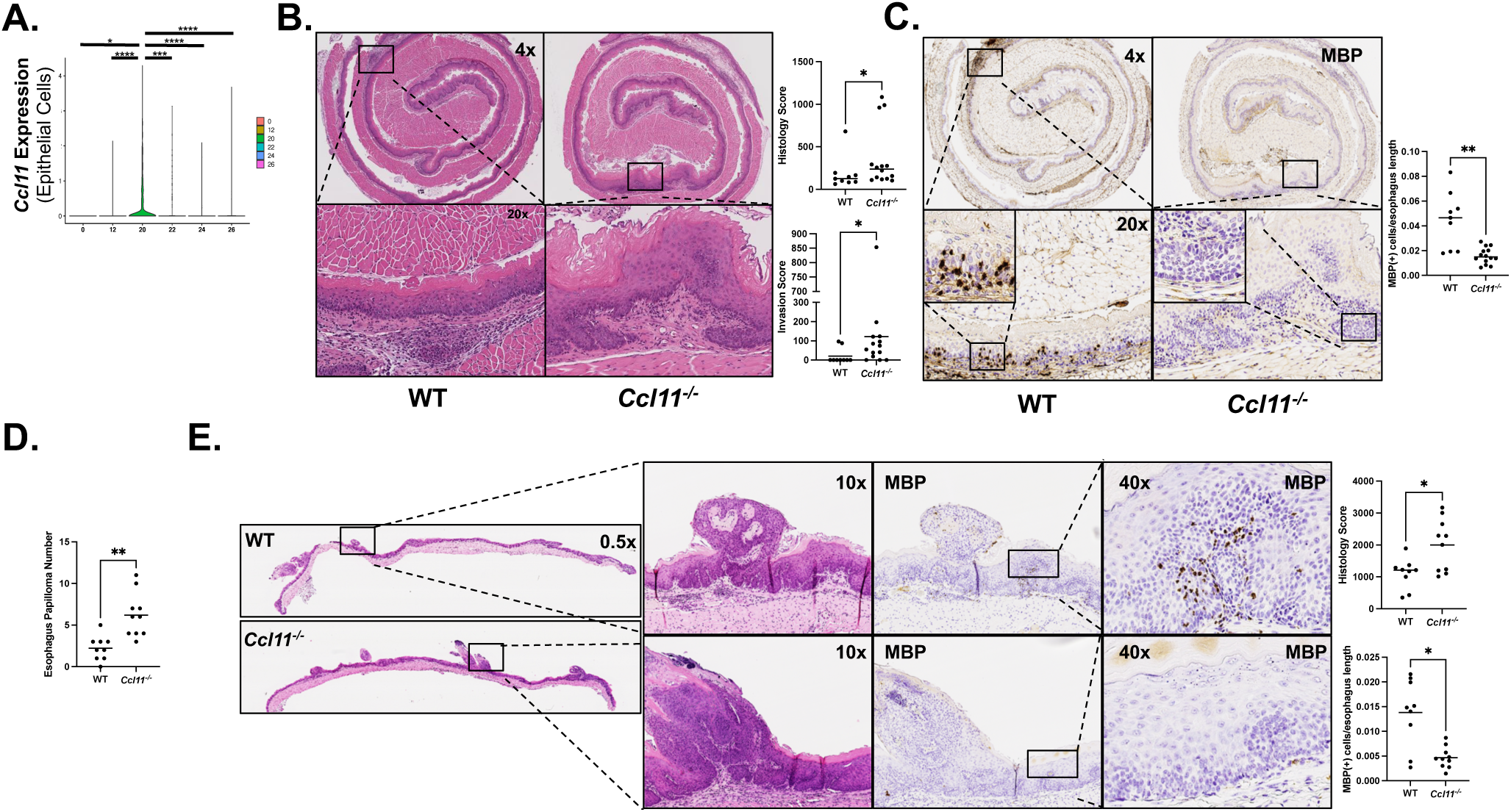
*Ccl11^-/-^*mice exhibit significantly worsened ESCC pre-cancer and carcinoma. (A) Analysis of sc-RNAseq (Yao *et al*., accession number CRA002118) shows that epithelial cell expression of *Ccl11* was significantly greater in what was termed hyperplasia (20 weeks after initiation of 4-NQO) as compared with 0 weeks (normal, P=0.01), 12 weeks (inflammation, P<0.001), 22 weeks (dysplasia, P=0.001), 24, weeks (carcinoma in situ, P<0.001), and 26 weeks (invasive carcinoma, P<0.001). Wilcoxon rank sum test was used to compare the target gene (i.e. *Ccl11*) expression between HYP and NOR, between HYP and INF, between HYP and DYS, between HYP and CIS and between HYP and ICA. Multiplicity was controlled with Bonferroni method. (B) *Ccl11^-/-^* mice display significantly increased histology score in the pre-cancer timeline (364.4 ± 95.6 vs 182.1± 63.9, P=0.04, n=9-14). They demonstrate significantly increased invasion parameter of the total score (121.8 ± 58.3 vs 20.5 ± 13.6, P=0.03, n=9-14). (C) *Ccl11^-/-^* mice had significantly fewer MBP positive cells/esophagus length as compared with WT controls (0.016 ± 0.0017 vs 0.044 ± 0.0076, P=0.002, n=9-14). (D) *Ccl11^-/-^* mice have significantly more esophageal papillomas than WT controls in a 4-NQO carcinoma timeline (6.1 ± 0.9 vs 2.2 ± 0.5, P<0.001, n=9-10). (E) *Ccl11^-/-^* mice demonstrate significantly greater histology score than WT controls in the same carcinoma timeline (1997 ± 257.8 vs 1104 ± 157.5, P=0.04, n=9-10). *Ccl11^-/-^* mice also show significantly fewer MBP positive cells/esophagus length than WT controls (0.0047 ± 0.00068 vs 0.014 ± 0.0023, P=0.02, n=9-10). For B-E, all comparisons are reported as mean ± SEM and are using Mann-Whitney.

### Ccl11^-/-^ mice exhibit worse tumorigenesis

Since *Ccl11* was upregulated at a timepoint when there were more eosinophils and downregulated when there were fewer eosinophils, we tested whether the absence of *Ccl11* would lead to worse neoplasia. *Ccl11^-/-^* mice showed significantly advanced disease after 8 week 4-NQO treatment (Figure 4B). Similar to the IL5mAb –treated and ΔdblGATA mice, the H&E highlights the greater degree of invasion in the *Ccl11^-/-^* mice. MBP IHC confirms that there were fewer eosinophils in *Ccl11^-/-^* mice (Figure 4C). After treatment with 4-NQO for 16 weeks, *Ccl11^-/-^* mice displayed significantly more esophageal papillomas (Figure 4D) and greater histology scores, (Figure 4E) again including increased invasion (Supplemental Figure 3). MBP IHC revealed significantly fewer eosinophils (Figure 4E). Just as in ΔdblGATA mice, *Ccl11^-/-^* mice displayed a trend towards decreased survival (Supplementary Figure 4A), and the experiment was concluded 4 weeks early as a consequence. However, *Ccl11^-/-^* mice also had a significantly greater number of tumors on the tongue (Supplemental Figure 4B) and thus it is difficult to interpret whether this change in survival was due to esophageal or oral tumorigenesis. Notably, while there were fewer eosinophils in the *Ccl11^-/-^* mice, it was not 0, which could be due to the presence of *Ccl24* or other unknown eosinophil chemoattractants.

### Recombinant IL-5 slows ESCC tumorigenesis

Since the reduction of eosinophil number exacerbated tumorigenesis, we next hypothesized that enhancing eosinophilic recruitment might reduce ESCC progression. In order to do so, we tested whether rIL-5 treatment, which increases eosinophilia, could slow neoplastic progression. We first treated mice weekly with intranasal rIL-5 in the second eight weeks of the pre-cancer protocol (Figure 1C). Mice treated with rIL-5 had significantly lower histology scores (Figure 5A) and significantly more MBP positive cells along the length of the esophagus (Figure 5B). The H&E shown highlights the increased smoothness of the epithelium, i.e. fewer rete pegs, a feature of pre-cancer. There was no difference in invasion score, but so few control mice develop invasion in this timeline it is not surprising that there was no difference. The same experiment was conducted in the carcinoma timeline, where WT mice were treated intranasally with rIL-5 weekly for the last 12 weeks of the experiment (i.e. after 4-NQO was stopped, see Figure 1C). Mice treated with rIL-5 had significantly fewer esophageal papillomas (Figure 5C) and less severe features of carcinoma, including invasion score (Figure 5D). There were significantly more MBP positive cells than controls (Figure 5E), confirming rIL-5 increases recruitment of eosinophils to the esophagus.

**Figure 5.**
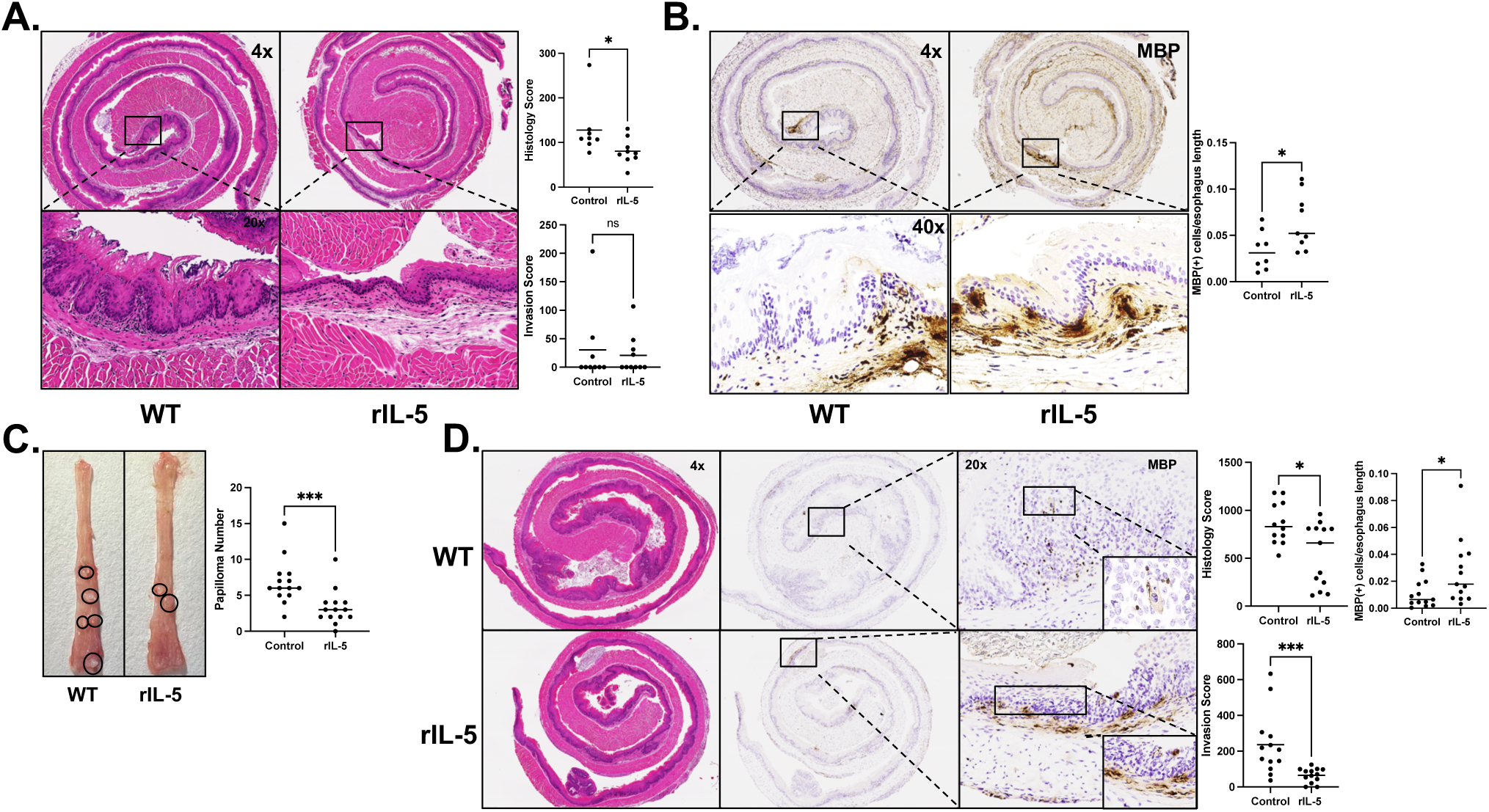
Intranasal recombinant IL-5 attenuates 4-NQO pre-cancer and carcinoma. (A) Mice treated with intranasal recombinant IL-5 (rIL-5) for the second eight weeks of the 4-NQO dysplasia timeline have significantly reduced histology score (80.5 ± 9.7 vs 127.8 ± 21.5, P=0.04, n=8-9), but there was no difference in the invasion parameter of the total score. (B) Intranasal rIL-5 results in significantly greater MBP positive cells/esophagus length as compared with WT control in the pre-cancer timeline (0.064 ± 0.010 vs 0.034 ± 0.0074, P=0.04, n=8-9). (C) Mice treated with intranasal rIL-5 during the final 12 weeks of the 4-NQO carcinoma timeline have significantly reduced esophageal papillomas (3.3 ± 0.6 vs 6.9 ± 0.8, P<0.001, n=14). (D) Mice treated with intranasal rIL-5 have significantly lower histology score than WT controls (537.8 ± 90.9 vs 863.9 ± 62.3, P=0.03, n=12-13), a lower invasion parameter of the total score (65.7 ± 11.1 vs 237.3 ± 48.8, P<0.001, n=12-13), and a significantly greater number of MBP positive cells/esophagus length (0.027 ± 0.0068 vs 0.010 ± 0.003, P=0.02, n=12-13). All comparisons are reported as mean ± SEM and are using Mann-Whitney.

### RNA-sequencing demonstrates eosinophilic contribution to oxidative state in esophageal squamous pre-cancer

Having established that the number of eosinophils modifies tumor progression, we then performed RNA-sequencing on the whole esophagus of WT and ΔdblGATA mice in the precancer model to understand how the presence of eosinophils affects the pre-cancer esophageal environment. After generating a list of differentially expressed genes between the two groups, we input this list into ingenuity pathway analysis (IPA) software to understand which molecular pathways were significantly different. Several of the most dysregulated pathways were related to metabolic alterations, and most of them were downregulated in the absence of eosinophils, including Oxidative phosphorylation, Glutathione Redox Reactions, Glutathione-mediated Detoxification, and eNOS signaling (Figure 6A). The list of all significantly dysregulated pathways is uploaded in GEO.

**Figure 6.**
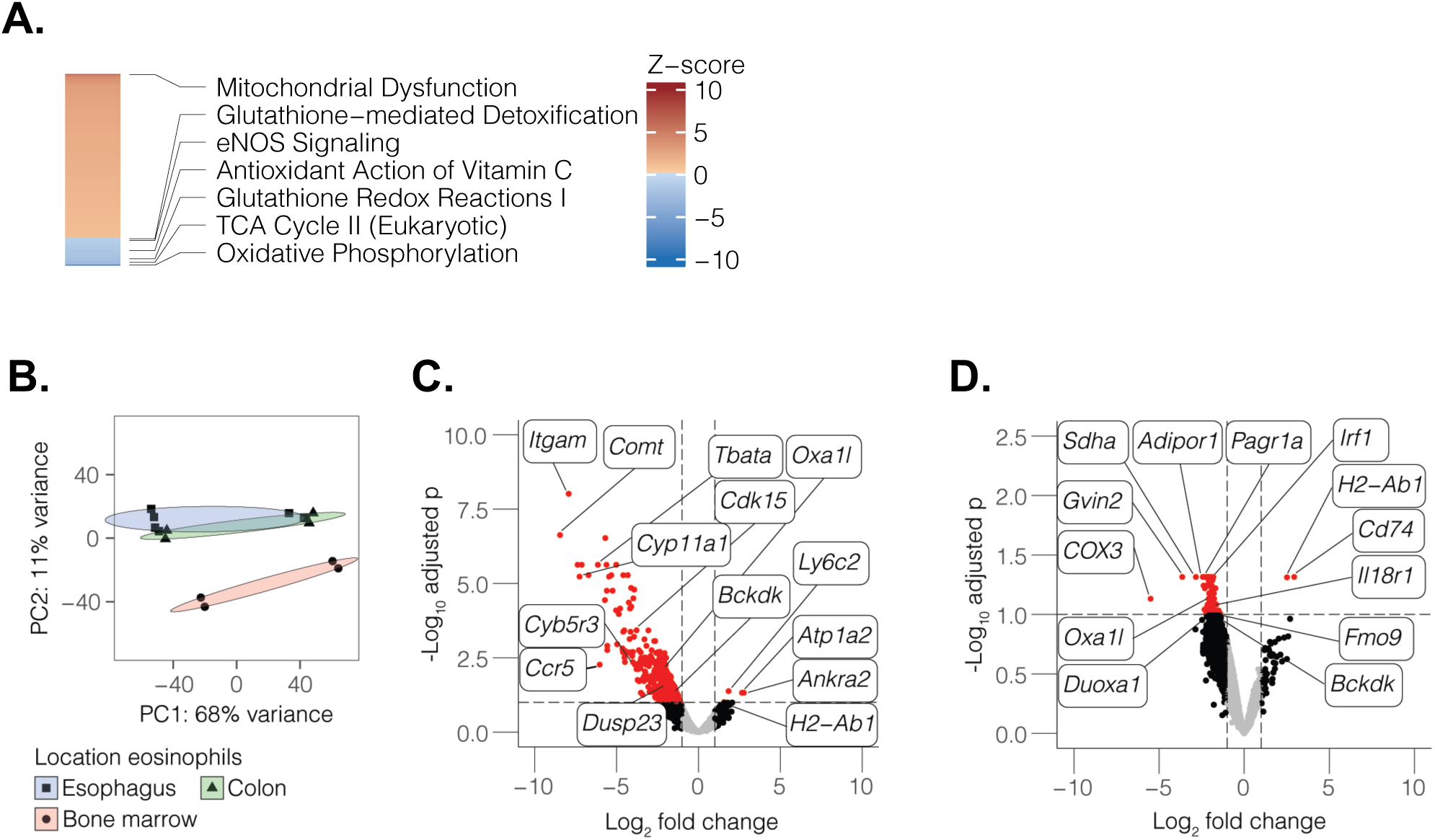
RNA-sequencing of tissue and eosinophils reveals a significant eosinophilic contribution to oxidative stress in esophageal squamous pre-cancer. (A) WT and ΔdblGATA mice (n=6-7) were treated with 4-NQO in the pre-cancer model, and after sacrifice the whole esophagus was collected for RNA. Bulk RNA-sequencing was performed. The list of differentially expressed genes was analyzed for pathway alterations using IPA, and a heatmap of z-scores for altered metabolic pathways is shown. Pathways that were statistically significant (adjusted p value <0.10) are shown and pathways of interest are annotated. The majority of oxidant pathways were significantly downregulated in the absence of eosinophils. (B) For eosinophil specific RNA-sequencing, eosinophils were retrieved using positive enrichment from the esophageal lamina propria (n=7) and from the colonic lamina propria from a random subset (n=4) of the same mice. Bone marrow was also extracted and eosinophils were induced *in vitro* from the same random subset (n=4). Eosinophils were subsequently subjected to bulk RNA-seq. Principle component analysis (PCA) of gene expression data is shown. (C) Volcano plot highlighting genes of interest in the comparison of esophageal eosinophils versus bone marrow eosinophils. Genes that are above the threshold for statistical significance and fold change are colored red. Many highlighted genes are mitochondria or directly involved in regulating oxidative stress, including *Sdha*, *Sdhd*, *Bckdk*, and *Oxa1l*. (D) Volcano plot highlighting genes of interest in the comparison of esophageal eosinophils versus colonic eosinophils. Again, many highlighted genes are mitochondrial or regulate oxidative stress, including *Sdha*, *COX3*, *Oxa1l*, *Duoxa1*, *Fmo9*, and *Bckdk*.

In order to study the transcriptional profiles of eosinophils specifically, we next isolated eosinophils from esophageal tissue of WT mice which had been treated with 4-NQO in the precancer model. Because eosinophils are not present in the esophagus normally, we isolated eosinophils from the colons of a random subset of the same mice. Additionally, we derived eosinophils from the bone marrow of a random subset of the same mice. We then compared the pre-cancer esophageal eosinophils to each of these groups. The principal component analysis (PCA) showed that the pre-cancer esophageal eosinophils were more similar to the colonic eosinophils than bone marrow-derived eosinophils (Figure 6B). When comparing esophageal precancer eosinophils with bone marrow-derived eosinophils (Figure 6C), we identified several differentially expressed genes which were either mitochondrial or related to redox state, including *Bco1*, *Cyb5r3*, *Cyp11a1*, *Sdha*, *Bckdk*, *Sdhd*, and *Oxa1l*. All of these genes were downregulated in the pre-cancer eosinophils. Similarly, when comparing pre-cancer esophageal eosinophils to colonic eosinophils, we again identified several genes related to redox state which were downregulated – *Sdha*, *Cox3*, *Oxa1l*, *Duoxa1*, *Fmo9*, and *Bckdk*. Notably, in this comparison, we also observed a significant upregulation of *Cd74*, the receptor of macrophage inhibitory factor (MIF), in the pre-cancer esophageal eosinophils. Additionally, *H2-Ab1* (part of MHCII) was upregulated in both comparisons in the pre-cancer esophageal eosinophils, suggesting antigen presentation by esophageal eosinophils may contribute to their function in attenuating esophageal cancer. The full list of dysregulated genes in each comparison is uploaded in GEO. Altogether, these results suggest pre-cancer esophageal eosinophils regulate other immune cells as well as metabolism.

### Apoptosis of pre-cancer and cancer cells is triggered by release of reactive oxygen species from degranulating eosinophils

Since pre-cancer eosinophils showed significant downregulation of several mitochondrial genes as compared with colonic and bone marrow-derived eosinophils, we hypothesized that precancer eosinophils may induce apoptosis of surrounding cells through release of reactive oxygen species (ROS) during degranulation. In order to test this, we performed Cleaved Caspase-3 and EPX immunofluorescent staining on randomly selected mice (n=4) treated with rIL-5 and 4-NQO for 16 weeks, controls treated with 4-NQO for 16 weeks (n=4), or ΔdblGATA mice treated with 4-NQO for 16 weeks (n=4). Mice treated with rIL-5 displayed a significantly greater number of CC3^+^ cells per HPF as compared with controls and ΔdblGATA mice (Figure 7A). Consistent with this, bulk RNA-seq from WT and ΔdblGATA mice with pre-cancer showed the apoptosis signaling pathway to be downregulated in the absence of eosinophils (Figure 7B). Moreover, immunofluorescence for EPX revealed that eosinophils were in close proximity to CC3 positive cells. Together, these experiments suggested that rIL-5 protects against the development of ESCC by inducing apoptosis of cells adjacent to eosinophils.

**Figure 7.**
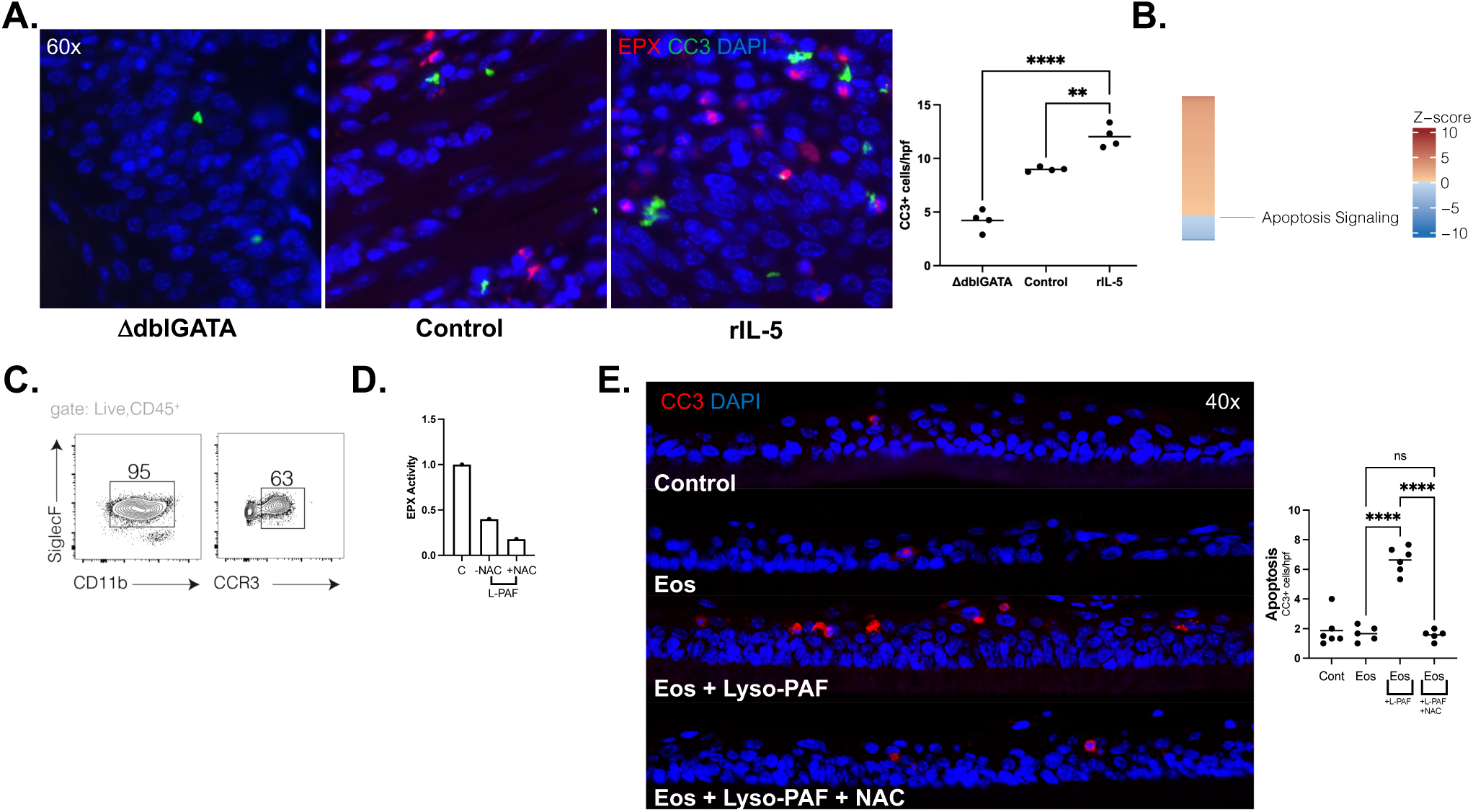
Eosinophils induce apoptosis of surrounding cells through release of ROS during degranulation. (A) Mice treated with intranasal rIL-5 have significantly more CC3 positive cells/high power field (hpf) as compared with WT controls (12.0 ± 0.5 vs 9.0 ± 0.1 CC3 positive cells/hpf, P=0.001, n=4 mice, 8-12 hpf/mouse) and ΔdblGATA mice (12.0 ± 0.5 vs 4.2 ± 0.49 CC3 positive cells/hpf, P<0.0001, n=4 mice, 8-12 hpf/mouse) treated with 4-NQO in the carcinoma timeline. A co-stain for EPX demonstrates close proximity of EPX positive cells with CC3 positive cells in both the rIL-5 and WT conditions. It also confirms the absence of eosinophils in the ΔdblGATA mice. (B) Heatmap of z-score of apoptosis pathway retrieved from IPA from tissue bulk-RNA sequencing from WT and ΔdblGATA mice (n=6-7), same as in Fig 6A. (C) This is an example of the flow cytometry gating strategy for bone marrow derived eosinophils. Eosinophils were defined as CD45+CD11b+Siglec-F+. They were further characterized by CCR3 status. (D) This is an example (one representative example from 3 different experiments) demonstrating that the addition of Lyso-PAF (L-PAF) results in degranulation of bone marrow eosinophils, as measured by an EPX activity assay. EPX activity induced by L-PAF was 40% of EPX activity induced by the detergent CHAPS, a positive control. N-acetylcysteine (NAC) reduced measured EPX activity from 40% to 18%. (E) Co-culture of bone marrow derived eosinophils with mouse precancer organoids grown in air-liquid interface shows that inducing degranulation of bone marrow eosinophils with L-PAF results in significantly greater CC3 positive cells as compared with control (6.6 ± 0.4 vs 1.9 ± 0.4, P<0.0001, n=2 independent experiments, 6 replicates), eosinophils alone (6.6 ± 0.4 vs 1.7 ± 0.2, P<0.0001, n=2 independent experiments, 5-6 replicates), or eosinophils plus L-PAF + NAC (6.6 ± 0.4 vs 1.6 ± 0.2, P<0.0001, n=2 independent experiments, 4-6 replicates). Images in (A) and (E) were taken using Keyence BZX-810. Images in (A) were done using a 60x oil immersion lens, and images in (E) were taken using a 40x lens.

Since rIL-5 treatment of 4-NQO mice demonstrated increased esophageal apoptosis we next sought to define the mechanism by co-culturing eosinophils with pre-cancer and cancer cells *in vitro*. To accomplish this, we cultured murine bone marrow derived eosinophils from WT mice, which were defined as CD45^+^CD11b^+^SiglecF^+^ cells [30]. Our culture technique resulted in 95% eosinophils (Figure 7C). These cells were then co-cultured with murine pre-cancer organoids which were grown to confluence on a transwell and then made three-dimensional by removing media from the apical side of the transwell, inducing differentiation. Eosinophils were cultured in the basolateral compartment with or without Lyso-platelet activating factor (Lyso-PAF), which induces eosinophil degranulation. Degranulation was confirmed by performing an Eosinophil Peroxidase assay (Figure 7D), which showed that Lyso-PAF resulted in 40% of degranulation as compared with the positive control CHAPS, similar to previous studies.[30] After 24 hours, the transwells were collected and fixed and immunofluorescence was done for CC3 to measure apoptosis. Without activation, these bone marrow-derived eosinophils did not induce increased apoptosis (Figure 7E, second image).

However, when degranulation was triggered with Lyso-PAF, there was significantly increased apoptosis of pre-cancer cells by CC3 immunofluorescence (Figure 7E, third image). Adding both Lyso-PAF and N-acetylcysteine, to buffer ROS, reduced the number of CC3 positive cells (Figure 7E, fourth image). This suggested that eosinophil degranulation was required for inducing apoptosis of surrounding cells and that the release of ROS during degranulation was at least one mechanism by which cell death occurred (Figure 7E).

We confirmed these results in a separate model using a human eosinophil cell line (EoL-1 cells) co-cultured with a human ESCC cell line (TE-11 cells). In these experiments, TE-11 cells were grown to confluence on transwells and then differentiated using an air-liquid interface model, similar to the mouse pre-cancer organoids. EoL-1 cells were differentiated in media with sodium butyrate in order to increase the percentage of mature eosinophils and cultured in the basolateral compartment of the transwell, similar to mouse bone marrow derived eosinophils. Differentiated EoL-1 cells were co-cultured with TE-11 cells with or without IL-5, a degranulating agent for human eosinophils. After 24 hours, transwells were fixed and immunofluorescent staining was performed for CC3 to assess apoptosis. When compared with control (i.e. co-culture with no EoL-1 cells), co-culture with differentiated EoL-1 cells minus IL-5 showed a trend towards greater apoptosis of TE-11 cells. However, adding IL-5 to differentiated EoL-1 cells led to significantly greater apoptosis of TE-11 cells as compared with control (Supplemental Figure 5), confirming our observation in the murine model.

Taken together, these data support that eosinophil degranulation is a driver of apoptosis of surrounding pre-cancer and cancer cells in our mouse model and that the release of ROS is one mechanism by which this occurs.

### Eosinophils suppress CD4 T cells in the pre-cancer esophageal squamous microenvironment

Having defined the direct effect of eosinophils on surrounding cells, we next sought to determine the indirect effect of eosinophils on other immune cells in the pre-cancer microenvironment. We performed IHC for CD4 T cells in WT, ΔdblGATA, and mice treated with rIL-5 in the pre-cancer model. There was a significantly greater number of CD4 T cells in the absence of eosinophils (Figure 8A) but significantly fewer in mice treated with rIL-5 (Figure 8B). Given this difference in CD4 T cells, we then performed a Luminex Multiplex Array on WT and ΔdblGATA mice to measure cytokines associated with T cell subsets. ΔdblGATA showed significantly increased IL-17A and a trend towards an increase in IL-6 and granulocyte-macrophage colony-stimulating factor (GM-CSF) (Figure 8C). The analytes for Th1, Th2, Th9, Th22, and Treg cells did not differ (Supplemental Figure 6). Bulk RNA-sequencing from WT and ΔdblGATA mice supported these data, as IL-6 signaling and IL-17 signaling pathways were enriched in ΔdblGATA mice in IPA (Figure 8D). Moreover, pathways downstream of IL-17, including EGF and NF-κB were also enriched in ΔdblGATA mice. Finally, the number of CD8 T cells was not different by IHC in the absence of eosinophils (Figure 8E) or with treatment of rIL-5 (Figure 8F). Thus, our data support that esophageal eosinophils might regulate Th17 function.

**Figure 8.**
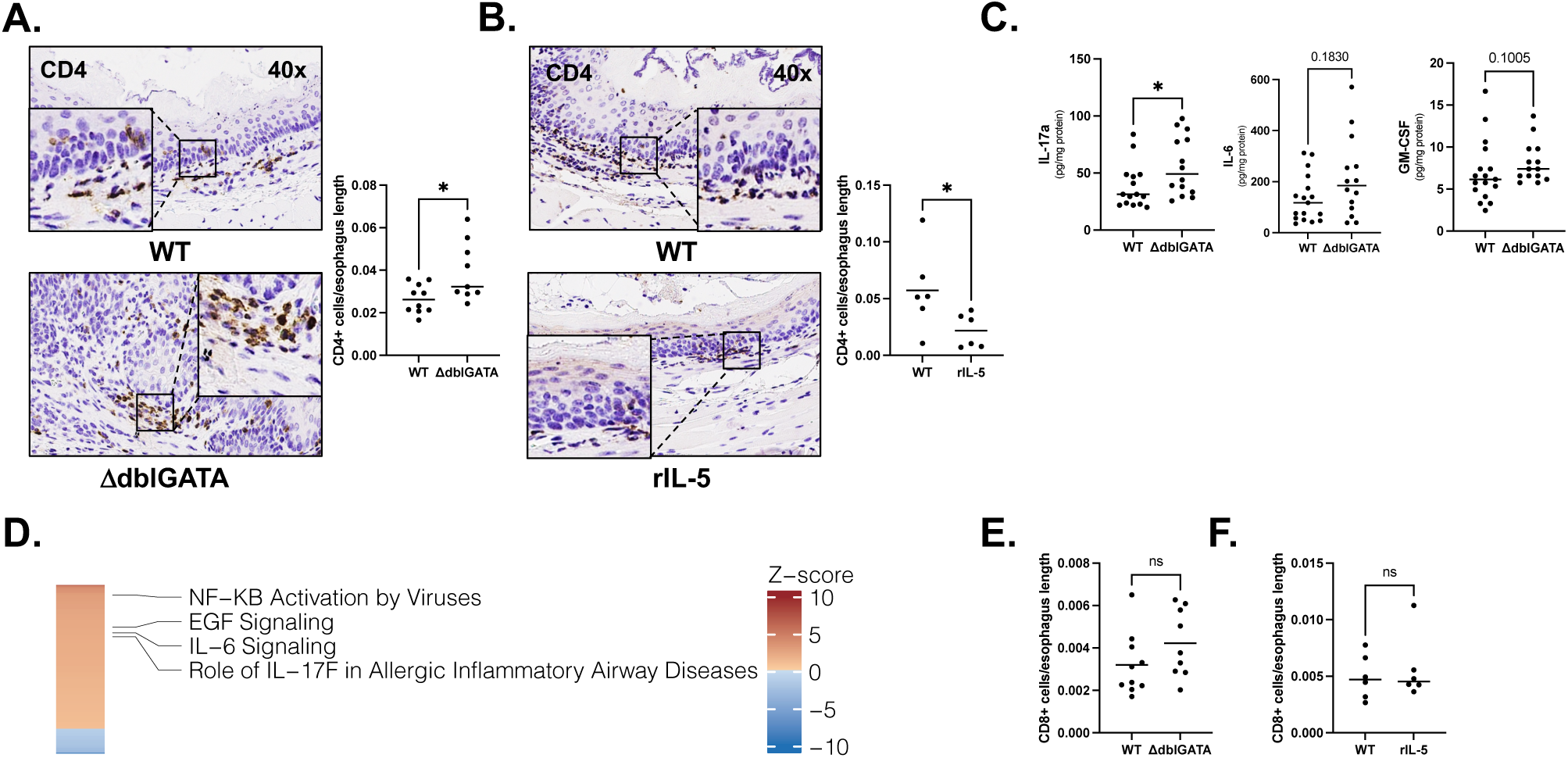
The absence of eosinophils alters the pre-cancer immune microenvironment and results in enrichment of pro-tumorigenic pathways. (A) ΔdblGATA have significantly more CD4 T cells/esophagus length as compared with WT mice (0.04 ± 0.005 vs 0.03 ± 0.002, P=0.04, n=9-10). (B) WT mice have significantly more CD4 T cells/esophagus length as compared with WT mice treated with rIL-5 (0.05 ± 0.01 vs 0.02 ± 0.006, P=0.02, n=6). (C) Luminex profiling shows IL-17a is significantly upregulated (56.6 ± 7.0 vs 38.3 ± 5.0 pg/mg protein, P=0.04, n=14-15) in ΔdblGATA mice. IL-6 (213.2 ± 41.9 vs 133.6 ± 22.5 pg/mg protein, P=0.18, n=14-17) and GM-CSF (8.2 ± 0.6 vs 7.0 ± 0.9 pg/mg protein, P=0.10, n=14-17) demonstrate a trend towards increase in ΔdblGATA mice. (D) Heatmap of z-scores of pathways retrieved from IPA from tissue bulk-RNA sequencing from WT and ΔdblGATA mice (n=6-7), same as in Fig 6A. IL-6 signaling, EGF signaling, Nf-κB signaling, and IL-17 are all enriched in ΔdblGATA mice. There is no difference in CD8+ cells/esophagus length in WT vs ΔdblGATA mice (E) or in WT vs rIL-5 treated WT mice (F). All comparisons are reported as mean ± SEM and are using Mann-Whitney.

## Discussion

While eosinophils are known effectors in allergic disease, new functions have been discovered in the pathobiology of cancer. [31] We investigated the role of eosinophils in ESCC and discovered that activated eosinophils are present in human ESCC, to a greater degree in early stage versus late stage cancers. Since tissue-resident eosinophils normally are not present in the esophagus, eosinophils are being specifically recruited during tumorigenesis. Similar to human ESCC, we then showed that there are increased eosinophils in pre-cancer as compared with cancer in the 4-NQO murine model of ESCC, suggesting that this model is relevant for studying eosinophil biology. We demonstrated that expression of *Ccl11* by epithelial cells and at some time points fibroblasts increases during the pre-cancer period when there are the most eosinophils, suggesting that CCL11 is one mechanism by which eosinophils are recruited in this model. We then demonstrated that reducing eosinophil number either pharmacologically through IL5mAb treatment or genetically with either eosinophil deficient mice or *Ccl11^-/-^* mice leads to worse precancer and carcinoma, suggesting eosinophils protect against the development of ESCC. Conversely, increased eosinophilia dramatically reduces the severity of pre-cancer and cancer, notably invasive disease. RNA-sequencing of whole esophageal tissue and eosinophils revealed that pre-cancer esophageal eosinophils have significantly reduced expression of several genes which regulate oxidative stress. Finally, our *in vitro* co-culture experiments collectively demonstrate that degranulation of eosinophils leads to increased apoptosis of surrounding precancer and cancer cells and that the release of ROS is one mechanism by which eosinophils can induce apoptosis.

To the best of our knowledge, this study is the first to define the contribution of eosinophils in ESCC. Specifically, it is the first to illustrate that eosinophils are activated in tissues from ESCC patients from the US and Japan using EPX IHC. It is also the first to demonstrate that treatment with intranasal rIL-5 can attenuate the number of tumors in the 4-NQO model, suggesting that this may be a future avenue for therapy in ESCC. Notably, there is one other recently published study which showed that mice treated with 4-NQO and ovalbumin, which induces a Th2 response that leads to increased eosinophilic infiltration, had reduced severity of ESCC as compared with mice treated with 4-NQO alone.[32] We were the first to show the direct cytotoxic effect of ROS on pre-cancer cells in a co-culture modeling system, though ROS have been previously implicated as an anti-tumorigenic mechanism. [33] It is also notable that in murine colon cancer activated eosinophils displayed an interferon signature,[12] but RNA-seq from eosinophils from esophageal pre-cancer showed significantly decreased *Irf1* as compared with colonic eosinophils and bone marrow derived eosinophils, suggesting this is not the mechanism of activation in murine esophageal squamous cancer. Additionally, the Luminex array showed that IFN-γ was trending towards being lower in Δdbl-GATA mice in the pre-cancer model. Since esophageal pre-cancer eosinophils had significantly upregulated *Cd74*, the receptor for MIF, a future direction is to more thoroughly investigate the role of MIF in ESCC development. This is one possible mechanism for how eosinophils are activated in esophageal squamous pre-cancer, as MIF has been shown to activate eosinophils in allergic disease.[34] Furthermore, based on the RNA-seq data, we hypothesize that antigen presentation by eosinophils contributes to reduced disease burden observed in our study.

Finally, while eosinophils have previously been shown to suppress Th17 differentiation in the small intestine, [35] we are the first to demonstrate that this may be occurring in the context of esophageal squamous malignancy. This is important because IL-17 has been shown to be protumorigenic in several mouse cancer models, including colon, lung, and skin, and inhibition of IL-17 has been shown to reduce metastasis.[36] Additionally, the enrichment of the IL-6, EGF, and NF-κB signaling in eosinophil-deficient mice, all pro-tumorigenic pathways downstream of IL-17[36], is supporting evidence eosinophils suppressing IL-17 and pro-tumorigenic effects. Our lab is currently further investigating the role between eosinophils and IL-17 in the 4-NQO model. In this vein, it is notable that numbers of CD8 T cells were not different in eosinophil-deficient mice or mice treated with rIL-5, as eosinophils have been shown to enhance CD8 T cells in melanoma.[37] In summation, for cancers in which eosinophils may be protective, this study provides evidence that strategies should be developed to increase recruitment of eosinophils to the tumor microenvironment.

In addition to the aforementioned new findings, another strength of our study is the use of multiple mouse models to demonstrate that reduction of eosinophil number leads to exacerbated tumorigenesis. We utilized 3-D coculturing methods to show the direct effect of eosinophils on pre-cancer and cancer cells. Eosinophils have been shown to have direct cytotoxic effect in other contexts as well, including in mastocytoma cells (P815 cells), [38] hepatocellular carcinoma cells (MH134 cells), [39] fibrosarcoma, [40] and melanoma (HBL and B16-F10 cells). [41]

With that being said, there are important limitations to consider as well. Notably, while CCL11 is an important chemoattractant for eosinophil recruitment, it also has been shown to have effects on epithelium. [42] Thus, it is possible that the exacerbation of disease in *Ccl11^-/-^* mice may be partially related to epithelial changes as well. Additionally, in our co-culture studies, murine eosinophils and EoL-1 cells have significant differences from human eosinophils. For example, neither makes all of the granules that human eosinophils do. Another limitation is that intranasal delivery of recombinant IL-5 likely results in eosinophil-mediated lung damage. As such, one of our future directions is to test whether different delivery methods of rIL-5 can also attenuate tumorigenesis.

From a patient-oriented perspective, tumor associated eosinophilia across all cancers has been associated with increased overall survival (though not disease-free survival),[18] suggesting that the presence of eosinophils could be a biomarker for a favorable prognosis. Since we have observed a greater number of EPX positive cells in earlier stage ESCC, this may be the case for this cancer as well. However, many of these studies utilize H&E to quantify eosinophil number instead of EPX or MBP IHC. Future studies should determine whether eosinophil number, perhaps by EPX or MBP IHC, predicts disease free survival in this cancer.

In conclusion, while eosinophils have context-dependent functions, we discovered that eosinophils are recruited to the early esophageal neoplastic environment at least in part by Ccl11 and protect from progression to ESCC. We determined that treatment with rIL-5, augmenting tissue eosinophilia, reduced tumorigenesis in an ESCC animal model. We showed that the mechanisms by which eosinophils are likely protective is through secretion of ROS and suppression of IL-17. Thus, therapies which either deliver more eosinophils to the esophagus or increase eosinophil recruitment to the esophagus should be further investigated in the treatment of ESCC.

## Materials and Methods

### ESCC patient samples

Our Japanese colleagues (MN) in Juntendo University constructed a tissue microarray (TMA) with paraffin embedded sections from 28 ESCC patients from the center of tumor, edge of the tumor, and tumor-adjacent normal tissue. Paraffin-embedded sections from 30 additional patients who underwent endoscopic submucosal dissection (ESD) as curative therapy were also provided.

At Vanderbilt University Medical Center, a pathology database was queried for ESCC patients from 2012-2020. Biopsies and resections were reviewed for availability of blocks. Of these, 14 resections and 32 biopsies were identified. One TMA was constructed from the resections and a second from biopsies. All patients were categorized by AJCC T stage (8^th^ edition).

### Human Eosinophil Peroxidase (EPX) Immunohistochemistry (IHC)

Detailed methods for staining and quantification are provided in Supplementary Methods. Slides were digitally scanned by the Vanderbilt Digital Histology Shared Resource (DHSR) core, which uses an Aperio AT2 and Leica SCN400 Slide Scanner for high-resolution bright field scanning. All slides were uploaded into QuPath [43] to analyze whole-slide images.

### Mice

All experiments were begun with 8-12-week-old C57BL/6J mice. These mice were maintained in controlled conditions with a consistent diet and kept in normal light/dark cycle, temperature, and humidity.

## 4-Nitroquinoline-1-oxide (4-NQO) treatment

Mice were given 4-NQO in two different timelines. To model pre-cancer, mice were given 4-NQO for 8 weeks followed by propylene glycol vehicle for 8 weeks. [22, 44, 45] For carcinoma modeling, mice were given 4-NQO for 16 weeks and then propylene glycol vehicle for 12 weeks. [20, 46, 47] Additional details are provided in Supplementary Methods.

### Histology scoring

Histology scoring was performed in accordance with previously published guidelines. [27] Further details are in Supplementary Methods.

### IL-5 Monoclonal Antibody Treatment

In the pre-cancer protocol, WT mice were treated with 4-NQO for the first 8 weeks of the experiment. For the second eight weeks, half the mice were given a 0.1 mg IL5mAb weekly (TRFK5, BioXCell) and the other half given 0.1 mg IgG1 isotype control (BioXCell) weekly by intraperitoneal injection. This dose reduces intestinal eosinophils. [12]

### Recombinant IL-5 Treatment

In the pre-cancer protocol, WT mice were treated with 4-NQO for the first 8 weeks of the experiment. For the second eight weeks, half the mice were given recombinant IL-5 (R&D) 100 ng weekly intranasally and the other half given PBS vehicle control. This dose was similar to what has been previously published to induce lung eosinophilia. [48] In the carcinoma protocol, the same dose was given for the final 12 weeks of the experiment.

### Single cell RNA sequencing (sc-RNAseq) analysis

The single-cell RNA sequencing data analyzed in this study are available from Yao *et al.* through GSA (Genome Sequence Archive in BIG Data Center, [29] Beijing Institute of Genomics, Chinese Academy of Sciences, http://gsa.big.ac.cn) under the accession number CRA002118. Visualization of target gene expression in each cell type was accomplished with Seurat v4.0. [49] Further details are in Supplementary Methods.

### Eosinophil and tissue RNA-sequencing (RNA-seq) and analysis

For details regarding eosinophil and tissue RNA-seq, please see Supplementary Methods.

### Bone marrow-derived eosinophils

Bone marrow eosinophils were isolated and differentiated using a previously published protocol [30] with modifications detailed in Supplementary Methods.

### Eosinophil degranulation EPX activity assay

Eosinophil degranulation was assessed *in vitro* in technical triplicates as previously described [50] with modifications detailed in Supplementary Methods.

### Immunofluorescent Staining

Fluorescent immunohistochemistry (IHC) on paraffin-embedded sections was performed as previously described. [51] Detailed methods are in Supplementary Methods.

### Statistics

Statistical analysis outside of RNA-seq and sc-RNAseq (see above) was performed in GraphPad Prism (v9.5.0). For two groups, the Mann-Whitney non-parametric test was used. For more than two groups, Kruskall-Wallis was used. Survival curve analysis was done using the Logrank (Mantel-Cox) test. Data are presented as mean ± standard error (SEM), except in analysis of human data, where they are presented as median with interquartile range (25%, 75%). For all experiments, the ROUT (Q=1%) test was performed to exclude outliers. P<0.05 was considered statistically significant.

### Study Approval

Human studies were approved by the Vanderbilt University Medical Center IRB (192400) and the Hospital Ethics Committee of Juntendo University Hospital (E21-060). All mouse experiments were approved by Vanderbilt University Medical Center Institute for Animal Care and Use Committee (V1900036-01) and Tennessee Valley Health System.

**All authors had access to the study data and have reviewed and approved the final manuscript**.

## Author contributions

Conception: YAC. Experimental design: YAC, CSW, JJ, LC. Acquisition of data: YAC, JJ, ZA, LS, JC, FR, AK, MN, TM, SM, HO. Analysis and interpretation of data: YAC, CSW, JJ, LC, JMP, MB, MW, SPS, MKW, GH, MKG, JAG. Intellectual contribution: all authors. Histopathological assessment: MKW, YAC. Single-cell RNA-seq analysis: LS and TK. Writing of manuscript: JJ and YAC. Critical revision of manuscript: all authors.

## Acknowledgements and Funding Sources

TE-11 cells were kindly provided by Anil Rustgi. The Vanderbilt Flow Cytometry Shared Resource, The Translational Pathology Shared Resource, Division of Animal Care core facilities all supported this work. We thank Richard M. Peek, Keith T. Wilson, Ariel Munitz, Elizabeth Jacobsen, and Marc Rothenberg for their critical advice in constructing the manuscript.

This work was supported by a VA-Career Development Award IK2BX004648, ACS-Institutional Research Grant, DDRC Pilot and Feasibility Grant Application P30 058404, a Vanderbilt Burroughs Welcome Supporting Careers in Research for Interventional Physicians and Surgeons (SCRIPS), and 5T32DK007673 (to YAC). It was supported by Veterans Affairs Merit Review grants I01BX004366 (to LAC) and I01BX001426 (CSW). It was supported by the National Institute of Diabetes and Digestive and Kidney Diseases (R03DK123489 to JAG, K01DK123495 to SPS, and 1K23DK13141 to GH). It was also supported by Japan Society for the promotion of Science (JSPS) Grant-in-Aid for Scientific Research (KAKENHI, 22K08878 to MN) and by Subsidies for Current Expenditures to Private Institutions of Higher Education from the promotion and Mutual Aid Corporation for Private Schools of Japan (to MN). Tissue morphology Core Services were performed through Vanderbilt University Medical Center’s Digestive Disease Research Center (P30DK058404 to MKW). It was supported by a VICC SPORE in GI cancer grant (P50CA236733 to MKW). We would also like to acknowledge the TPSR (supported by NCI/NIH Cancer Center Support Grant 5P30CA68485-19) for assistance with immunohistochemistry (Shared Instrumentation Grant S10 OD023475-01A1). We would like to thank the Digital Histology Shared Resource (DHSR) for assistance with data storage and processing.

## Writing Assistance

No individuals provided writing assistance besides the authors listed.

## Data Transparency

The single-cell RNA sequence data analyzed in this study are available from Yao et al. through GSA (Genome Sequence Archive in BIG Data Center, Beijing Institute of Genomics, Chinese Academy of Sciences, http://gsa.big.ac.cn) under the accession number CRA002118. Tissue and eosinophil RNA-seq from mice have been deposited in GEO under the accession numbers GSE233448 and GSE233443. Further details on how to access these files is in the supplement. Supporting analytic code can be made available upon request.

