## Supplementary Materials for "Eosinophils exert direct and indirect anti-tumorigenic effects in the development of esophageal squamous cell carcinoma"

Human Eosinophil Peroxidase (EPX) Immunohistochemistry (IHC)

Esophageal squamous cancer paraffin blocks and tissue microarrays were cut at 5 microns on positively charged slides. The slides were de-paraffinized in three changes of xylene and hydrated to water in a series of graded alcohols. The slides underwent antigen retrieval using a citrate buffer (pH 6.0) solution at 105°C in a pressure cooker for 20 minutes then cooled at room temperature for 10 minutes. The slides were washed in distilled water and placed in Tris-buffered saline with 0.1% Tween (TBST) wash buffer solution prior to continuation of the staining protocol. Endogenous enzymes were blocked using a 0.03% (H202) peroxidase block solution for 5 minutes, rinsed in wash buffer, and then incubated for one hour in the primary antibody (1:1000 primary dilution, EPX Mayo Clinic, clone MM25.82.2.1). The slides were gently rinsed in wash buffer and a peroxidase labelled polymer was applied using Dako EnVision + System-Horseradish Peroxidase labeled polymer for 30 minutes. After a gentle rinse again in wash buffer the slides were treated with a DAB+ Substrate-Chromogen for 5 minutes to complete the staining protocol. The slides were washed in distilled water, counterstained in Mayer's Hematoxylin, blued in running tap water and dehydrated in 3 changes of absolute alcohol and cleared in xylene prior to cover slipping.

To calculate the percentage of total cells which were EPX positive, automated cell detection for both total cells and EPX positive cells was utilized in QuPath to account for differences in size of TMA cores and ESD specimens. In all TMAs, each patient had 2-4 cores per patient, and EPX positive cells/total cells were averaged for all cores.

Mice

*Ccl11*$^{-/-}$ mice were obtained from Lori Coburn and have been previously described. [1] The breeding strategy for these mice was *Ccl11*$^{+/-}$ x *Ccl11*$^{+/-}$. ΔdblGATA mice were obtained from Jackson Laboratory (Strain #005653). The breeding strategy for these mice was $X^{WT}Y$ (wild-type males) x $X^{Mut}X^{WT}$ (heterozygote females) and $X^{Mut}Y$ (knockout males) crossed with $X^{Mut}X^{WT}$ (heterozygote females) to generate wild type and knockout mice from the same parents. WT (wild-type) mice were obtained from Jackson Laboratory.

4-NQO Treatment

4-NQO (Acros 203792500) was dissolved in propylene glycol (Fisher Scientific P355-4) vehicle as a stock solution at a dosage of 50 mg/mL. A new stock solution was prepared weekly and stored at 4°C. The 4-NQO stock solution was then further diluted in drinking water of mice at a dosage of 100 μg/mL, which is the same dose reported in several other publications.[2-4] Water with 4-NQO was replaced weekly. Mice were continuously allowed access to drinking water.

In the experiment in which mice in the carcinoma protocol were compared with those in the pre-cancer timeline, all three experimental groups were coordinated to begin and end at the same time. The 8 week 4-NQO treatment group was given only vehicle for the first 12 weeks of the experiment, then received 4-NQO treatment for 8 weeks, and then given only vehicle for the last 8 weeks of the experiment. The 16 week 4-NQO treatment group was given 4-NQO for 16 weeks and then given only vehicle for the last 12 weeks of the experiment.

For all experiments, each cage contained WT and knockout mice in order to decrease cage effect. In IL5MAB and rIL-5 experiments, experimental and control treatments were equally distributed among mice in the same cage to decrease the impact of cage effects. For all experiments, the bedding was mixed amongst all cages with experimental mice monthly.

Allocation and conduct of each experiment (i.e. weighing mice weekly, changing bedding) in each experiment was done by either ZA or AK, who were blinded to genotype.

After sacrifice, the esophagus was removed from all mice, the inside of the esophagus washed with phosphate-buffered saline (PBS) with a gavage needle and splayed longitudinally. Tumors in the esophagus were counted under a dissecting microscope. Counting was done by an investigator who was blinded to genotype (MAB, ZA, or YAC). Then, the esophagus was rolled with the distal esophagus in the center and the proximal esophagus on the outside of the roll. Then, the Swiss-roll was fixed in 10% buffered formalin for 24 hours, transferred to 70% ethanol overnight, and then embedded in paraffin. 5 μm cross-sections of each sample were cut and stained with hematoxylin and eosin (H&E) for histologic analysis. Paraffin embedding, sectioning, and staining was performed by the Vanderbilt Tissue Pathology Shared Resource (TPSR). For the *Ccl11*$^{-/-}$ carcinoma experiment, the esophagus was fixed after being splayed longitudinally and sections were cut along the longitudinal axis. Additionally, since 4-NQO can cause oral tumors, the tongue was examined in all experiments and the number of tongue tumors was recorded.

Histology Scoring

Slides were digitally scanned by the Vanderbilt Digital Histology Shared Resource (DHSR) core. All slides were uploaded into QuPath, a free open source software, to analyze whole-slide images. Briefly, four parameters which are most commonly observed in mice treated with 4-NQO were used to determine severity of histology score: rete pegs, nuclear histologic abnormalities, papillomas, and invasive carcinoma. Rete pegs and nuclear histologic abnormalities are the most common features in mice treated with 4-NQO for 8 weeks (i.e. during pre-cancer), while papillomas and invasive carcinoma are more common in mice treated with 4-NQO for 16 weeks (i.e. during cancer). Severity and degree of each parameter was determined using QuPath. These parameters were scored by an expert GI pathologist who was blinded to mouse genotypes (MKW). In the results section, we report both the total histology score and highlight the invasion parameter of the total score (labeled in the graphs as "invasion score") as it represents the most severe feature of disease.

Major Basic Protein (MBP) IHC

Slides were placed on the Leica Bond Max IHC stainer. All steps besides dehydration, clearing and coverslipping were performed on the Bond Max. Slides were deparaffinized. Enzymatic induced antigen retrieval was performed on the Bond Max using Proteinase K (Dako, Agilent, Santa Clara, CA) for 5 minutes. Slides were incubated with anti-MBP (1:1000, Lee Laboratory, Mayo Clinic) for one hour and then incubated in a rabbit anti-rat secondary antibody (1:2000, BA-4001, Vector Laboratories, Inc.) for 15mins. The Bond Polymer Refine detection system was used for visualization. Slides were then dehydrated, cleared and coverslipped. This IHC was performed by our Vanderbilt TPSR.

Major Basic Protein (MBP) quantification was done in Qupath by highlighting the region of interest and using the positive cell detection feature with a consistent threshold. In the first experiment, where pre-cancer eosinophilia was compared with eosinophilia in papillomas and invasive carcinoma, the region of interest was defined by presence of pre-cancer features (i.e. rete pegs, nuclear histologic abnormality) versus presence of papilloma or invasive carcinoma. This was then standardized to the length of that region of interest, as measured using the polyline tool. For experiments with IL5MAB, ΔdblGATA, *Ccl11*$^{-/-}$, or recombinant IL-5, the major purpose of MBP quantification was to illustrate that there were lesser or greater eosinophilic infiltration. Thus, we standardized the MBP quantification to the entire esophagus length, as sections of swiss rolls can vary slightly.

### Single Cell RNA sequencing (sc-RNAseq) analysis

A time-ordered single-cell transcriptomic profiling was conducted on six esophageal lesions. Mice were sacrificed before (week 0), during (week 12), and after treatment (weeks 20, 22, 24, or 26), and were categorized into six stages: normal (NOR), inflammation (INF), hyperplasia (HYP), dysplasia (DYS), carcinoma in situ (CIS), and invasive carcinoma (ICA). These data include $CD45^+$ immune cells [29975 (cells) by 14283 (genes)] and $CD45^-$ non-immune cells [36114 (cells) by 15596 (genes)]. Immune cells were categorized into T cells, B cells, myeloid cells, and natural killer cells; and non-immune cells were categorized into epithelial cells, fibroblasts, endothelial cells, and myocytes.

Wilcoxon rank sum test was used to compare the target gene (i.e. *Ccl11*) expression between HYP and NOR, between HYP and INF, between HYP and DYS, between HYP and CIS and between HYP and ICA in each cell type. Multiplicity was controlled with Bonferroni method, and adjusted p-values $\leq 0.05$ were used for statistical significance. All statistical analyses were conducted using R version 4.1.

Example scripts to process and analyze data are available at https://github.com/ ~. Detailed code will be available from the corresponding authors upon request.

### Acquisition of lamina propria cells

Cells from the colonic lamina propria were acquired as previously described [5] with the following modifications to enhance viability of eosinophils: (1) 10% FBS instead of 5% FBS was used in incubation steps that include serum; (2) The digestion media consisted of 10 mL HBSS with calcium and magnesium instead of RPMI 1640; (3) The digestion media contained 1.5 mg/mL collagenase and no Liberase; (4) The concentration of DNAse was increased to 0.10% DNAse I (Sigma-Aldrich, D50250); (5) The maximum centrifugation speed was reduced to 370 x *g*, except for the density gradient (475 x *g*). For cells from the esophageal lamina propria, the esophagus was flushed with PBS, opened longitudinally, and cut into 4 equal-sized pieces. The esophageal tissue was then subjected to the same protocol as the colonic lamina propria with the following modifications to increase eosinophil yield: (1) The initial volume of incubation with DTT was reduced to 5 mL; (2) the volume of digestion media was reduced to 10 mL and the collagenase concentration was 1.0 mg/mL.

Selection of lamina propria eosinophils

Eosinophils were acquired using positive immunomagnetic selection with the siglecF$^+$ selection kit (Miltenyi Biotec, Gaithersburg, MD) according to the manufacturers' instruction with the following modifications: (1) Buffer was supplemented to contain a final concentration of 5 mM EDTA and 0.05% DNAse was added; (2) The working volume was 15 μL buffer with 5 μL of beads. Flow cytometry was used to confirm purity of eosinophils.

RNA isolation

Extraction of RNA from whole tissue was performed on tissue stored in RNA-later (Sigma-Aldrich)until homogenization using a Tissue-Tearor (Dremel, Racine, WI), followed by phenol/chloroform extraction as described [6] and clean-up with the Rneasy Mini Kit with on-column DNAse digestion according to manufacturers' instruction. For eosinophil RNA-seq, cells were processed using the Ovation RNA-seq System V2 (Tecan, Mannëdorf, Switzerland) by the Vanderbilt Technologies for Advanced Genomics core.

RNA-sequencing

Quality control for RNA was performed by the Vanderbilt Technologies for Advanced Genomics core using RNA 6000 Pico (Agilent, Santa Clara CA). For eosinophil RNA-seq the cDNA library was prepared using a NEB library preparation kit. Paired end 150bp sequencing was performed on a NovaSeq 6000 (Illumina, San Diego, CA).

RNA-sequencing analysis

For tissue bulk RNA-seq, analysis was performed as previously described.[5] For eosinophil RNA-seq, limma was used for differential expression analysis. The tissue of origin for eosinophils, i.e. esophagus, colon, or bone marrow, was used as metadata in the linear model. Cage or which mouse the cells originated from did not measurably affect gene expression as assessed by consensus correlation coefficient (not shown).

Differential expression analysis

For differential expression analysis, limma (version 3.50.3) was used on log-CPM transformed counts with prior count set to 3 or DESEq2 (version 1.34.0) was used on non-normalized counts. Unless otherwise indicated, genes with an adjusted p value <0.05 ($-\log_{10}$ of 0.05 is 1.3) and $\log_2$ fold change >|1| were considered differentially regulated. For heatmaps with gene counts, a Z-score, i.e. the number of standard deviations above or below the mean, of normalized counts was calculated using scale function in R. Ingenuity pathway analysis (IPA) was used for pathway assessment. Genes with an adjusted p value of <0.05 were used as input. Annotation was done with AnnotationDbi (version 1.56.2) using org.Mm.eg.db (version 3.14.0). Images were generated with pheatmap (version 1.0.12), complexheatmap, RColorBrewer (version 1.1-3), ggplot2 (version 3.3.5), and EnhancedVolcano (version 1.12.0).

Bone Marrow Derived Eosinophils

Bone marrow cells were recovered from the tibiae and femurs of WT C57BL/6J mice by spinning individual bones within a cut-off 200 uL pipette tip placed in a 1.5 mL eppendorf at 10,000 x *g* for 10 minutes at room temperature in FACS buffer (PBS with 2% FBS and 2 mM EDTA) on day 0. Red blood cell lysis was done twice with ACK lysis buffer (ThermoFisher). Cells were plated at $1x10^6$ cells/mL in complete RPMI1640 supplemented with 105 ng/mL mouse FLT3 (Biolegend) and 100 ng/mL mouse SCF (Biolegend). Cells were washed and re-plated on day 4, 8, 10, and 12 at the original volume in complete RPMI supplemented with 10 ng/mL recombinant mouse IL-5. Only day 4 were cells were added back to the original plate. On other days, new plates were used. Cells were acquired on day 14 and the percentage of eosinophils was determined by flow cytometry with mouse $CD4^+$ splenocytes used as negative control.

Eosinophil degranulation EPX activity assay

Eosinophils were mycoplasma free as assessed by a Mycoplasma detection kit (Applied Biological Materials, Vancouver, Canada). Eosinophils were resuspended in eosinophil degranulation media (RPMI1640 without phenol red) and plated in an untreated flat-bottom tissue culture plate. After resting at 37°C for 1 hour, secretagogues prepared in eosinophil degranulation media were added followed by a 30-minute incubation at 5% $CO_2$ at 37°C to degranulate eosinophils. Then, one volume of freshly prepared OPD buffer [0.5 mg/mL OPD (ThermoFisher) in 0.05 M citric acid, 0.05 M sodium phosphate with 0.09% $H_2O_2$ (v/v) (Sigma-Aldrich)] was added. Following incubation for 2-8 minutes, the reaction was stopped with 1 volume of 1M $H_2SO_4$. The plate was read at 490 nM with a GloMax® Discover Microplate Reader (Promega, Madison, Wisconsin). EPX activity values were corrected for with a media only control (blank). As a positive control, 100% EPX activity was defined by degranulation after treatment with CHAPS detergent (Thermofisher). EPX activity was measured after treatment of eosinophils with 35μM platelet activating factor (Lyso-PAF C16, Cayman Chemicals, Ann Arbor, Michigan, 60906) and 1 mM *N*-acetyl-L-cysteine (NAC, Sigma-Aldrich) in combination with PAF.

Co-culture of eosinophils with pre-cancer organoids

To derive mouse pre-cancer organoids, a WT mouse was treated with 4-NQO for 8 weeks followed by propylene glycol vehicle for 8 weeks. At that time, the mouse was sacrificed and esophagus removed. The esophagus was placed in 1 mg/mL dispase II (Roche) in a thermomixer at 37°C for 10 minutes. Under a dissecting scope, the epithelium was peeled off the submucosa and placed in 1 mL 0.25% trypsin-EDTA. This was incubated again in a thermomixer at 37°C for 10 minutes. The sample was vortexed for 10 seconds. The trypsin-EDTA was then removed and placed in 10 mL of soybean trypsin inhibitor (1 mg/mL, dissolved in HBSS, Thermofisher). The remaining tissue was then incubated for a second time in 1 mL fresh 0.25% trypsin-EDTA in a thermomixer at 37°C for 10 minutes. Again, the sample was vortexed for 10 seconds and then transferred to the 10 mL of soybean trypsin inhibitor. The sample (including media and tissue) was then filtered through a 40 μM strainer and rinsed with PBS. The filtered liquid was then centrifuged for 5 minutes at 1000 RPM and resuspended in organoid media. After counting, cells were again centrifuged at 1000 RPM and suspended in Matrigel (Corning, 356231) at 3000 cells per 20 μL plug. After Matrigel polymerized at 37°C for 20 minutes, organoid media was placed on top of Matrigel plug. Organoid media consisted of Advanced DMEM/F12 (12634010, Gibco), 1X B-27™ Supplement (17504044, Gibco), 1X GlutaMAX™ (35050061, Gibco), 1X N-2™ Supplement (17502048, Gibco), 1 mM HEPES (15630080, Gibco) and 2% (v/v) penicillin/streptomycin (15140122, Gibco)] supplemented with 20% (v/v) R-spondin-conditioned media (from R-spondin-expressing cells gifted by Dr. Jeff Whitsett, The University of Cincinnati, Cincinnati, OH) and 10% (v/v) Noggin-conditioned media (from Noggin-expressing cells gifted by Dr. G.R. van den Brink, Amsterdam, NL), EGF (500 μg/mL), and N-acetylcysteine (0.5 M). For initial plating and at the time of splitting organoids, Y27632 (10 μM, Thermofisher) was also added.

For co-culture with bone marrow-derived eosinophils, pre-cancer organoids were collected on day 3 after plating. Cells were incubated in trypsin-EDTA 0.25% with Y27632 10 μM and DNase (Sigma, 1:200) for 1 hour in order to achieve a single cell suspension. Then, 150,000 cells were plated on 0.4 μm transwells (Corning 3413). After 3 days, organoid media was removed from the apical side in order to create an air-liquid interface (ALI). After another 3 days, 500,000 eosinophils were placed into the basolateral compartment of the transwell with or without Lyso-PAF 35μM or N-acetylcysteine 1 mM. After 24 hours, transwells were collected, fixed in 10% neutral buffered formalin for 24 hours, switched to 70% ethanol, and submitted to Vanderbilt TPSR for processing, embedding, and cutting.

Co-culture of Eol-1 cells with TE-11 Cells

Eol-1 cells (Sigma, 94042252) were maintained as undifferentiated in RPMI media supplemented with 10% FBS and 1% Pen/Strep. Prior to co-culture, Eol-1 cells were differentiated in RPMI media with above supplements and 500 μM sodium butyrate for 6 days in order to increase the percentage of mature eosinophils.

Meanwhile, TE-11 cells, a generous gift from Anil Rustgi, were also cultured in RPMI supplemented with 10% FBS and 1% Pen/Strep. 75,000 TE-11 cells were plated per transwell for each experiment. Cells were grown to confluence on a transwell for 5 days, with apical and basolateral media changes every other day. Media was then removed from the apical side in order to create air-liquid interface (ALI) and the cells were allowed to differentiate for 5 days, changing the media in the basolateral compartment every other day. After 10 days, 500,000 differentiated Eol-1 cells were placed into basolateral compartment for co-culture with TE-11 cells in ALI. Eol-1 cells were co-cultured with TE-11 cells with or without 10 ng/mL recombinant IL-5. After 24 hours, the transwell membranes were cut out, fixed in 10% neutral buffered formalin for 24 hours, switched to 70% ethanol, and submitted to Vanderbilt TPSR for processing, embedding, and cutting.

Flow cytometry

For cell surface staining, cells were incubated in antibody cocktail for 20 minutes at 4°C in the dark. Samples were blocked using 30 μL normal rat serum (StemCell Technologies, Vancouver, Canada). Flow cytometric analysis was performed using a 4-Laser Fortessa or 5-laser LSRII (BD, San Jose, California) with FACSDiva software (BD). Analyses were performed using FlowJo (BD). For all flow experiments, a live/dead stain (ThermoFisher, Waltham, Massachusetts) was used to only assess live cells. Antibodies used for flow cytometry are listed in Supplementary Table 1.

Immunofluorescent Staining

Slides were deparaffinized using Citrosolve, rehydrated by graded alcohols, and permeabilized using TBS with 0.05% Tween-20. Antigen retrieval consisted of 10-minute boiling in 10mM pH6 sodium citrate. After blocking, samples were incubated with the following primary antibodies overnight at 4°C: anti-cleaved caspase 3 (Cell Signaling Technologies, 1:200), anti-β-catenin (BD Biosciences, 1610153, 1:500), or anti-EPX Clone MM25.82.2.1 (Mayo Clinic, 1:250). Secondary antibodies were conjugated to either 488 or 594 Alexafluor dyes (1:500, Thermo Fisher Scientific) and incubated for 2 hours at room temperature. Nuclear staining was performed using ProLong Gold antifade reagent with DAPI (Thermo Fisher Scientific). Fluorescent stained slides were visualized on a Keyence BZ-X810 upright microscope. Cleaved caspase 3 (CC3) positive cells were quantified per high power field using a 40x objective, but images shown in Figure 6 were taken using a 60x oil immersion lens to highlight the important features of the image. 8-12 high power fields were averaged per mouse, and data is represented as high power field per mouse.

Luminex Multiplex Array

Luminex Multiplex Array was performed as previously described. [7] Esophageal tissues were harvested at the time of sacrifice and homogenized using a handheld pestle-type rotary homogenizer in CelLytic MT lysis extraction reagent from Sigma-Alrich (St. Louis, MO, USA). The 17 analyte ThermoFisher Mouse Cytokine Th1/Th2/Th9/Th17/Th22/Treg 17-Plex Mouse ProctaPlex Panel was performed according to the manufacturer's instructions and analyzed on a FLEXMAP 3D instrument (Luminex, Austin, TX, USA). Tissue protein concentrations, as assessed by the bicinchoninic acid (BCA) protein assay kit (Pierce, Rockford, IL, USA) were used to standardize the analyte data.

Statistics

As mentioned in the main text, a ROUT test (Q=1%) was performed to exclude outliers. Outliers were present in some experiments based on this test: recombinant IL-5 pre-cancer (n=1 in each group) and recombinant IL-5 carcinoma (n=1 in control group). There were no outliers by this test in other experiments.

**Supplementary Tables**

Supplementary Table 1 – List of FACS antibodies

| Antigen-label | Manufacturer | Catalog number |
|---|---|---|
| CD45-BV785 | Biolegend | 103149 |
| Siglec-F-BV421 | BD | 562681 |
| CD11b-PerCP/Cy5.5 | eBioscience | 45-0112 |
| CCR3-FITC | R&D Systems | FAB729F |

Supplementary Table 2 – Patient Demographic Information

| Case Number | Sex | Age | Stage | Surgical Resection, ESD, EMR, or Biopsy | Prior Treatment |
|---|---|---|---|---|---|
| 1 | M | 73 | T1a | Resection | None |
| 2 | M | 77 | T1b | Resection | None |
| 3 | M | 70 | T1b | Resection | None |
| 4 | M | 73 | T3 | Resection | None |
| 5 | F | 79 | T2 | Resection | None |
| 6 | F | 66 | T3 | Resection | None |
| 7 | F | 71 | T1b | Resection | None |
| 8 | M | 61 | T1b | Resection | None |
| 9 | M | 68 | T1b | Resection | None |
| 10 | F | 62 | T2 | Resection | None |
| 11 | M | 63 | T1b | Resection | None |
| 12 | F | 61 | T3 | Resection | None |
| 14 | M | 61 | T3 | Resection | None |
| 15 | M | 60 | T3 | Resection | None |
| 16 | M | 52 | T1b | Resection | None |
| 17 | F | 81 | T1b | Resection | None |
| 19 | M | 73 | T1b | Resection | None |
| 20 | F | 75 | T1b | Resection | None |
| 21 | M | 63 | T1b | Resection | None |
| 22 | M | 74 | T1a | Resection | None |
| 23 | F | 49 | T2 | Resection | None |
| 24 | M | 76 | T3 | Resection | None |
| 25 | F | 67 | T1a | Resection | None |
| 26 | M | 82 | T3 | Resection | None |
| 27 | M | 78 | T1a | Resection | None |
| 28 | M | 60 | T1b | Resection | None |
| 29 | M | 75 | T1b | Resection | None |
| 30 | M | 77 | T3 | Resection | None |
| 31 | M | 77 | T1a | ESD | None |
| 32 | M | 78 | T1a | ESD | None |

| | | | | | |
|---|---|---|---|---|---|
| 33 | M | 77 | T1a | ESD | None |
| 34 | F | 62 | T1a | ESD | None |
| 35 | M | 77 | T1a | ESD | None |
| 36 | M | 60 | T1a | ESD | None |
| 37 | M | 77 | T1a | ESD | None |
| 38 | M | 83 | T1a | ESD | None |
| 39 | M | 50 | T1a | ESD | None |
| 40 | M | 70 | T1a | ESD | None |
| 41 | M | 77 | T1a | ESD | None |
| 42 | M | 74 | T1a | ESD | None |
| 43 | M | 78 | T1a | ESD | None |
| 44 | M | 77 | T1b | ESD | None |
| 45 | M | 64 | T1a | ESD | None |
| 46 | M | 70 | T1a | ESD | None |
| 47 | M | 73 | T1a | ESD | None |
| 48 | M | 76 | T1a | ESD | None |
| 49 | F | 65 | T1a | ESD | None |
| 50 | M | 57 | T1a | ESD | None |
| 51 | M | 89 | T1a | ESD | None |
| 52 | M | 60 | T1b | ESD | None |
| 53 | F | 72 | T1a | ESD | None |
| 54 | M | 77 | T1a | ESD | None |
| 55 | M | 70 | T1a | ESD | None |
| 56 | M | 62 | T1b | ESD | None |
| 57 | F | 79 | T1a | ESD | None |
| 58 | M | 53 | T1a | ESD | None |
| 59 | F | 76 | T1a | ESD | None |
| 60 | F | 66 | T1a | ESD | None |
| 61 | M | 71 | T1a | Resection | Chemotherapy and radiation |
| 62 | M | 54 | T3 | Resection | Chemotherapy and Radiation |
| 63 | M | 61 | T3 | Biopsy | Chemotherapy and Radiation |
| 64 | M | 63 | T1a | Resection | Chemotherapy and Radiation |
| 65 | M | 76 | T1b | EMR | Chemotherapy and Radiation |
| 66 | M | 89 | Tis | Biopsy | None |
| 67 | M | 48 | T4a | Resection | Chemotherapy and Radiation |
| 68 | F | 43 | Tis | Resection | None |
| 69 | M | 51 | T1a | EMR | None |
| 70 | M | 72 | T2 | Resection | Chemotherapy and Radiation |
| 71 | M | 44 | T1b | Resection | None |
| 72 | M | 53 | T4a | Resection | Chemotherapy and Radiation |

| | | | | | |
|---|---|---|---|---|---|
| 73 | M | 74 | T3 | Resection | Chemotherapy and Radiation |
| 74 | M | 62 | T3 | Resection | Chemotherapy and Radiation |
| 75 | M | 47 | T3 | Biopsy | None |
| 76 | M | 75 | T3 | Biopsy | None |
| 77 | M | 50 | T4 | Biopsy | None |
| 78 | M | 47 | T3 | Biopsy | None |
| 79 | M | 83 | T4 | Biopsy | None |
| 80 | F | 63 | T4 | Biopsy | None |
| 81 | M | 56 | T4 | Biopsy | None |
| 82 | M | 71 | T3 | Biopsy | None |
| 83 | M | 56 | T4 | Biopsy | None |
| 84 | M | 57 | Tis | Biopsy | None |
| 85 | F | 52 | T3 | Biopsy | None |
| 86 | M | 55 | T4 | Biopsy | None |
| 87 | M | 76 | T3 | Biopsy | None |
| 88 | F | 90 | T3 | Biopsy | None |
| 89 | M | 54 | T4 | Biopsy | None |
| 90 | M | 66 | Tis | Biopsy | None |
| 91 | F | 73 | Tis | Biopsy | None |
| 92 | M | 69 | T4 | Biopsy | None |
| 93 | M | 59 | T4 | Biopsy | None |
| 94 | F | 74 | Tis | Biopsy | RFA |
| 95 | F | 70 | T2 | Biopsy | None |
| 96 | F | 65 | T3 | Biopsy | None |
| 97 | M | 59 | T1a | Biopsy | Chemotherapy and radiation |
| 98 | F | 56 | T3 | Biopsy | None |
| 99 | M | 81 | T4 | Biopsy | None |
| 100 | F | 63 | T3 | Biopsy | None |
| 101 | M | 66 | T4 | Biopsy | None |
| 102 | F | 68 | Tis | Biopsy | None |
| 103 | M | 76 | T3 | Biopsy | None |
| 104 | F | 59 | T3 | Biopsy | None |
| 105 | M | 67 | T3 | Biopsy | None |
| 106 | M | 83 | Tis | Biopsy | None |

Supplementary Table 3 - Comparison of demographic information between patients with ≤T2 ESCC versus patients with ≥T3 ESCC

| | ≤T2 | ≥T3 | p-value |
|---|---|---|---|
| Age | 71 (62,77) | 62.5 (55.8, 74.3) | 0.02 |
| Gender (Female) | 18/66 (27%) | 9/38 (24%) | 0.81 |
| Prior Therapy | 6/66 (9%) | 6/38 (16%) | 0.35 |

### Supplemental Figures

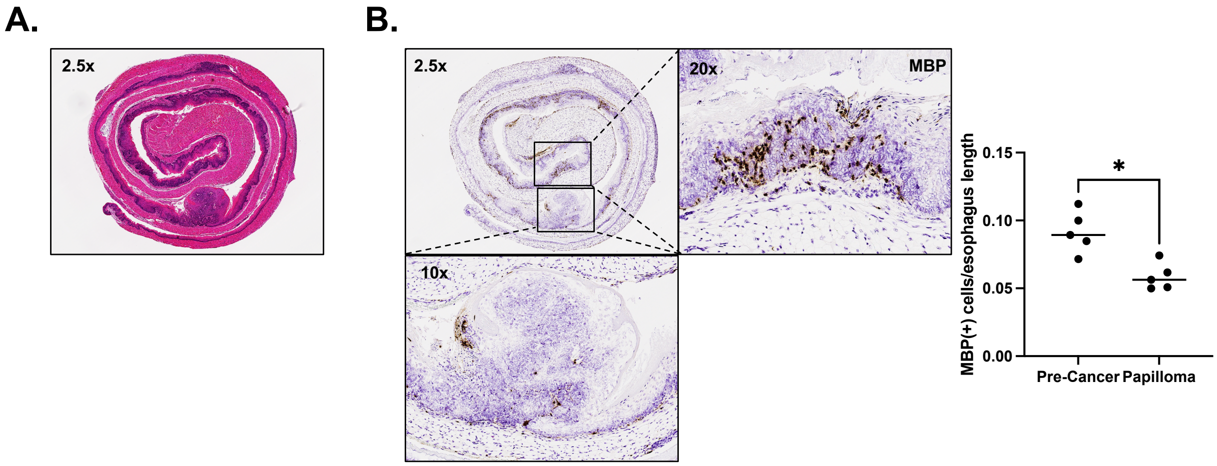


Supplemental Figure 1. Papillomas have greater numbers of eosinophils as compared with pre-cancer in the same mice. (A) Representative H&E of a WT mouse treated with 4-NQO in the carcinoma timeline. (B) Pre-cancerous areas show significantly greater MBP positive cells as compared with papillomas within the same mice (0.092 ± 0.0069 vs 0.059 ± 0.0044, P=0.02, n=4). MBP quantification was standardized to the length of pre-cancer or papilloma. All comparisons are reported as mean ± SEM and are using Mann-Whitney.

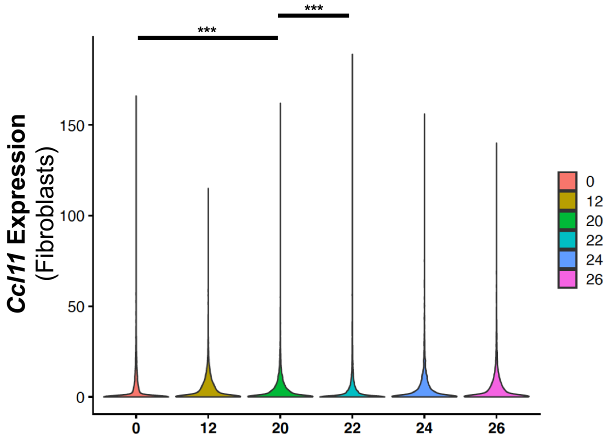


Supplemental Figure 2. 4-NQO increases *Ccl11* expression in fibroblasts during pre-cancer. Analysis of sc-RNAseq (Yao *et al*., accession number CRA002118) shows that fibroblast cell *Ccl11* expression was significantly greater in what was termed hyperplasia (20 weeks after initiation of 4-NQO) as compared with 0 weeks (normal, P<0.001) and 22 weeks (dysplasia, P<0.001). There was no difference between 20 weeks and 12 weeks (inflammation), 24 weeks (carcinoma in situ), or 26 weeks (invasive carcinoma). Wilcoxon rank sum test was used to compare the target gene (i.e. *Ccl11*) expression between HYP and NOR, between HYP and INF, between HYP and DYS, between HYP and CIS and between HYP and ICA. Multiplicity was controlled with Bonferroni method.

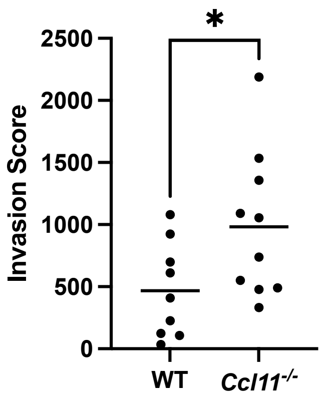


Supplemental Figure 3. *Ccl11*$^{-/-}$ mice have increased invasion score in carcinoma. *Ccl11*$^{-/-}$ mice have significantly increased invasion parameter of the total histology score as compared with WT (982.2 ± 185.0 vs 468.2 ± 126.6, P=0.04, n=9-10). This comparison is reported as mean ± SEM using Mann-Whitney.

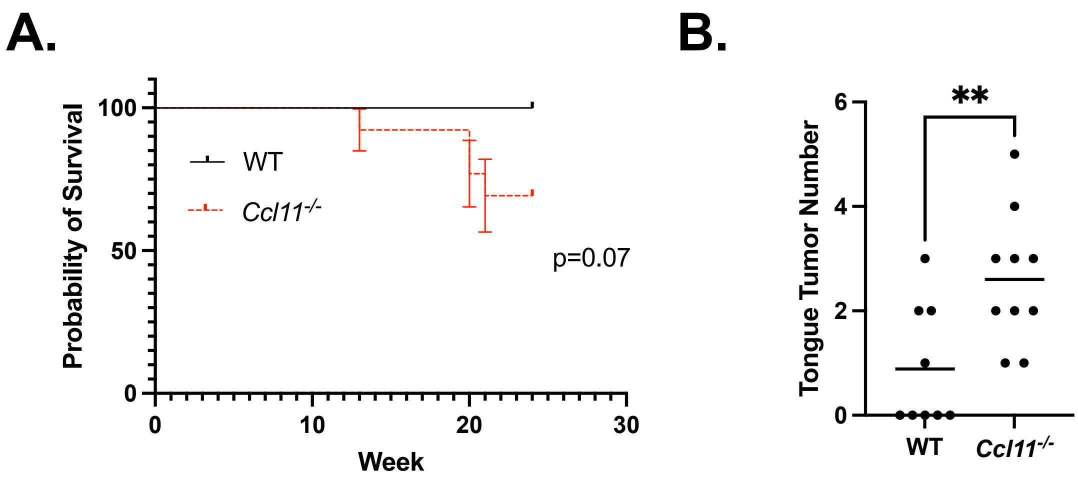


Supplemental Figure 4. *Ccl11*$^{-/-}$ mice have exacerbated oral tumorigenesis in the 4-NQO carcinoma timeline. (A) There is a trend towards worse survival in *Ccl11*$^{-/-}$ mice as compared with WT controls (P=0.07, n=9-14). (B) *Ccl11*$^{-/-}$ mice have significantly more tongue tumors as compared with WT controls (2.6 ± 0.4 vs 0.9 ± 0.4, P=0.009, n=9-10). Log-rank (Mantel-Cox) test was used for survival analysis, and tumor number is reported as mean ± SEM using Mann-Whitney

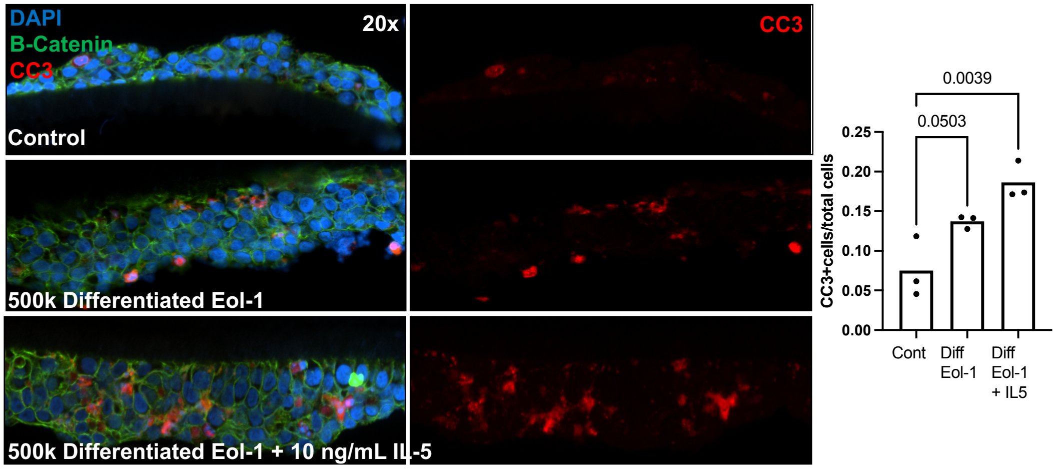


Supplemental Figure 5. Eol-1 cells co-cultured with an ESCC cell line results in apoptosis. Differentiated Eol-1 cells co-cultured with TE-11 cells grown in air-liquid interface results in a trend towards greater apoptosis as measured by CC3 positive cells/total cells as compared with control (0.14 ± 0.005 vs 0.075 ± 0.022, P=0.05, n=3 independent experiments). Differentiated Eol-1 cells with IL-5 co-cultured with TE-11 cells results in significantly greater apoptosis also measured by CC3 positive cells/total cells (0.19 ± 0.014 vs 0.075 ± 0.022, P=0.004, n=3 independent experiments). This comparison is reported as mean ± SEM using Kruskall-Wallis test.

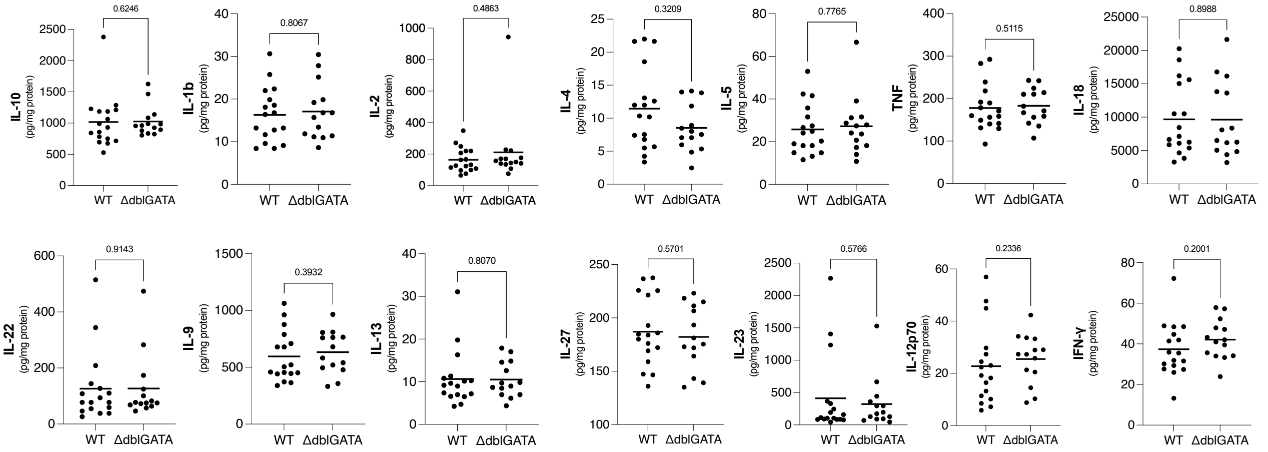


Supplemental Figure 6. Luminex profiling results for cytokines not significantly different between WT and ΔdblGATA mice in the pre-cancer timeline.
